# Lumped Parameter Liver Simulation to Predict Acute Hemodynamic Alterations Following Partial Resections

**DOI:** 10.1101/2022.12.27.522018

**Authors:** Jeffrey Tithof, Timothy L. Pruett, Joseph Sushil Rao

**Affiliations:** Department of Mechanical Engineering, University of Minnesota, Minneapolis, MN, USA; Division of Solid Organ Transplantation, Department of Surgery, University of Minnesota, Minneapolis, MN, USA; Schulze Diabetes Institute, Department of Surgery, University of Minnesota, Minneapolis, MN, USA

**Keywords:** Hepatic blood flow, hemodynamics, lumped parameter model, liver resection

## Abstract

Partial liver resections are routinely performed in living donor liver transplantation and to debulk tumors in liver malignancies, but surgical decisions on vessel reconstruction for adequate inflow and outflow are challenging. Pre-operative evaluation is often limited to radiological imaging, which fails to account for post-resection hemodynamic alterations. Substantial evidence suggests post-surgical increase in local volume flow rate enhances shear stress, signaling hepatic regeneration, but excessive shear stress has been postulated to result in small for size syndrome and liver failure. Predicting hemodynamic alterations throughout the liver is particularly challenging due to the dendritic architecture the vasculature, spanning several orders of magnitude in diameter. Therefore, we developed a mathematical lumped parameter model with realistic heterogeneities capturing inflow/outflow of the human liver to simulate acute perfusion alterations following surgical resection. Our model is parameterized using clinical measurements, relies on a single free parameter, and accurately captures established perfusion characteristics. We quantify acute changes in volume flow rate, flow speed, and wall shear stress following variable, realistic liver resections and make comparisons to the intact liver. Our numerical model runs in minutes and can be adapted to patient-specific anatomy, providing a novel computational tool aimed at assisting pre- and intra-operative surgical decisions for liver resections.

## Introduction

The liver receives dual blood supply from the portal vein and hepatic artery. The portal vein provides 75 – 80% of the blood flow draining from the splanchnic system, while the hepatic artery provides the remaining 20 – 25% of oxygenated blood (1). The two vascular systems branch independently across numerous generations throughout the liver, eventually reaching a confluence at the sinusoidal capillary channels. The blood from the liver sinusoids drains into a network of successive generations of merging centrilobular veins that form segmental hepatic veins that ultimately drain through the hepatic veins (left, middle, and right) into the inferior vena cava just below the diaphragm (2). The circulation of blood from the portal vein and hepatic artery provides a dual blood supply into well-demarcated liver segments, but blood exits the liver from three veins draining from shared territorial segments, making it a complicated and unique system for surgical resections (Fig. 1A).

**Fig. 1:**
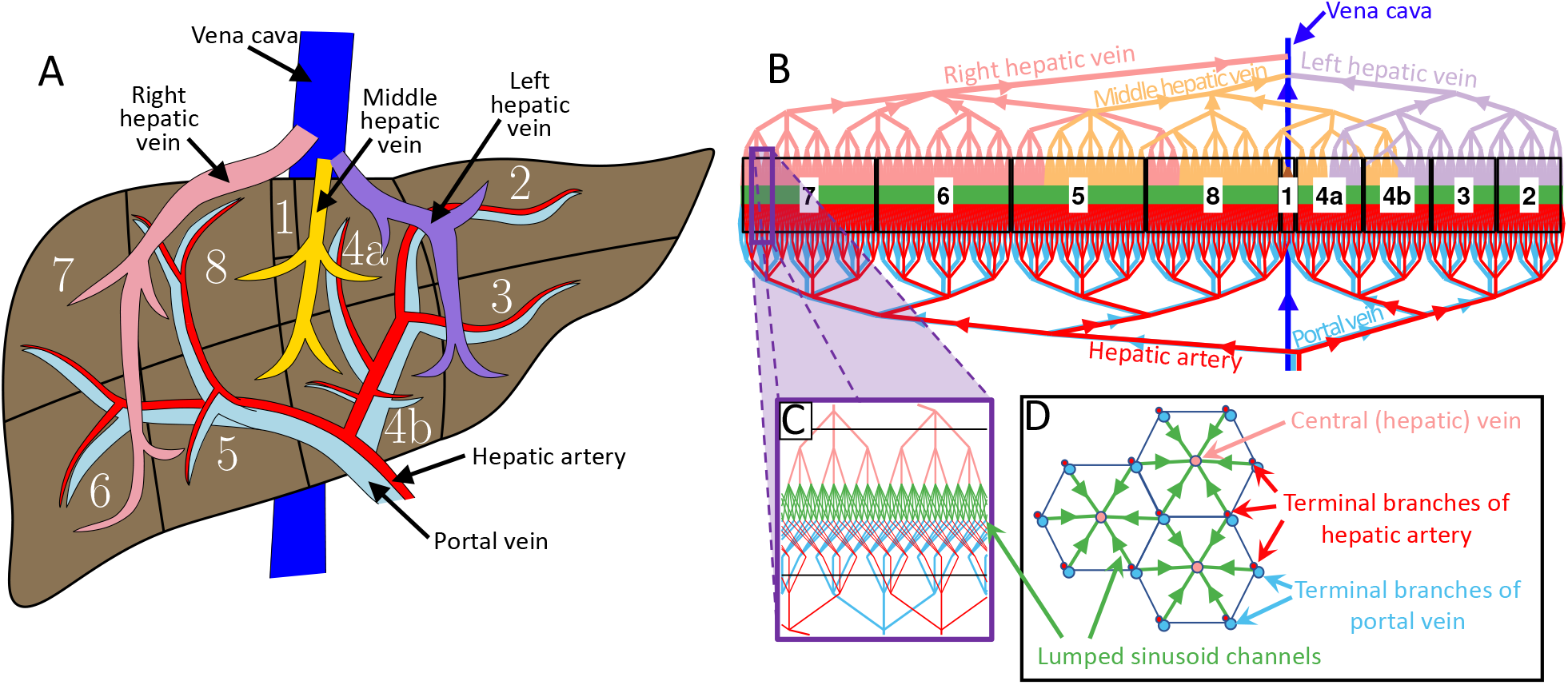
Schematic of the realistic model of human liver blood flow. (A) An illustration of liver anatomy, highlighting the eight segments and inflow/outflow vasculature for each liver segment. (B) The numerical model developed in this study represents liver segments, labeled with white numbered boxes, arranged linearly. Box size is proportional to liver segment volume fraction (indicated in Table 1). The hepatic artery, portal vein, sinusoids, vena cava, and three hepatic veins are color-coded, as indicated; these colors are used consistently throughout this article. The outflow of blood from liver segment 1 which often drains directly into the vena cava is plotted in brown and is barely visible. (C) An enlarged view of the idealized geometry of the sinusoidal microvasculature, which is obtained by considering (D) a segment of an infinite two-dimensional tiling of hexagonal lobules. Parallel branches of the hepatic artery and portal vein merge at the start of the sinusoids, with blood flowing toward a central (hepatic) vein. Each hepatic artery - portal vein is assumed to drain into three adjacent central veins, and each central vein receives blood from six adjacent hepatic artery/portal veins as is captured in our idealized network model (C). Thus, each lobule has six lumped sinusoid channels in our model. The network shown in (B) models a liver with only 730 lobules for the sake of visual simplicity and clarity.

**Table 1:** Parameters associated with liver segments used in this study. Liver volume fractions correspond to mean values reported by Abdalla et al. (15). Outflow fractions are estimates based on our prior clinical experience.

| Parameter | Liver segment |  |  |  |  |  |  |  |  |
| --- | --- | --- | --- | --- | --- | --- | --- | --- | --- |
|  | 1 | 2 | 3 | 4a | 4b | 5 | 6 | 7 | 8 |
| Liver volume fraction | 1.8 | 8.0 | 8.0 | 8.3 | 16.4 | 6.4 | 16.4 | 16.4 | 16.4 |
| Outflow fraction to LHV | 0 | 1 | 1 | 0.5 | 0.5 | 0 | 0 | 0 | 0 |
| Outflow fraction to MHV | 0 | 0 | 0 | 0.5 | 0.5 | 0.75 | 0 | 0 | 0.75 |
| Outflow fraction to RHV | 0 | 0 | 0 | 0 | 0 | 0.25 | 1 | 1 | 0.25 |
| Outflow fraction directly to vena cava | 1 | 0 | 0 | 0 | 0 | 0 | 0 | 0 | 0 |

Throughout all generations of inflow vasculature, the arrangement of a portal vein, hepatic artery, and bile duct is termed the “portal triad”. Six portal triads define the external boundaries of a compact hexagonal arrangement of parenchymal and non-parenchymal cells forming the functional unit of the liver called a “lobule” (3). The centrilobular vein at the cross-sectional center of each hexagon receives blood from the sinusoids that are formed by complex interconnections of terminal branches of a portal vein and hepatic artery (4). The sinusoids are lined by liver endothelial cells that respond to hemodynamic alterations with high sensitivity and transmit signals to the underlying parenchymal and non-parenchymal cells (5). (Note that “hemodynamic alterations” in this article is referring to hepatic hemodynamics and not systemic hemodynamics). Growing evidence supports this mechanism as the general signaling pathway initiating liver regeneration following surgical interventions. However, the liver has varying capacity for tolerance and regeneration following extensive resection. Beyond this threshold, extensive liver resection has resulted in hepatic insufficiency and has been deemed the main cause of mortality (6).

Liver resections are routinely performed for tumor debulking (7). Surgical techniques involve removal of one or more segments confined to specific hepatic territory of portal vein and hepatic artery branches. These procedures aim to downsize diseased liver volume and induce hypertrophy of the remnant liver to restore original liver volume. Staged hepatectomies, involving two or more surgical interventions, typically consist of an initial vessel ligation followed by liver resection (8). However, ligation of a primary branch of portal vein supplying a desired segment – without parenchymal transection – has been well described to not result in atrophy of the targeted regions, due to compensatory flow arising from the presence of intersegmental interconnections and collaterals (9). For any liver resection that includes parenchymal transection, the intricate, dendritic nature of the liver vasculature leads to heterogeneous alterations to perfusion which vary depending on the particular segment(s) resected, volume of remnant liver, and details of surgical reconstruction of the vasculature. Therefore, disruption of the portal network channels in the parenchyma not only results in hemodynamic alterations causing atrophy of the desired segment, but also initiates mechanical bio-mediation promoting regeneration (10).

Various liver resections involving a single segment or a combination of segments have been defined in the literature (11). Liver resections involving segments 4a, 4b, 5, and 8 are complex due to a common venous outflow through the middle hepatic vein (MHV). Often MHV resection is recommended and leads to congestion due to outflow obstruction, mandating MHV reconstruction (12–14). Inadequate pre- and intra-operative assessment of hemodynamic alterations may result in poor post-operative hepatic/metabolic function. Despite significant progress in surgical and radiological techniques (15–21), the association between resected volume and hemodynamic alterations in the remnant liver or transplanted graft remains unclear.

To predict these alterations to blood flow within the liver following resections, we developed a lumped-parameter model of blood flow through the entire human liver (Fig. 1B-D). Our general approach, which has been used previously to model hepatic blood flow (22–24) and liver resection (25–27), is based on a hydraulic analog of Ohm’s law, detailed below. Although many simulation approaches are possible (see the review article by Verma et al. (28)), lumped modeling is beneficial due to its computational tractability, enabling simulations of massive networks consisting of millions of lobules (27). The primary limitation is that the geometry is often overly simplified to facilitate straightforward generation of such massive networks (e.g., all branching is idealized as perfect bifurcations). The key novelties of our model are the incorporation of realistic heterogeneities to the vascular branching, demarcation of the eight liver segments via a spatially extended physical representation of the liver, and clinical measurements of deceased donor livers to provide model parameters. In the next section, we provide the full details of our computational model and the clinical measurements we performed to parameterize the model.

## Methods

### Lumped-parameter model with spatial structure for quantifying perfusion variations

The liver model described herein is implemented as a lumped parameter hydraulic network with fixed inlet and outlet pressures driving flow through a network of branching vessels with prescribed connectivity and variation in vessel diameter and length. We refer to each point at which a vessel branches or merges as a “node,” and we track the spatial coordinates and connectivity of the nodes. As depicted in Fig. 1B, the model is constructed with sequential generations plotted vertically, from the inflow of parallel right and left hepatic arteries (red) and portal veins (light blue) at the bottom, to the sinusoids (green) in the middle, and finally outflow through the right, middle, and left hepatic veins (abbreviated RHV, MHV, and LHV, plotted in pink, yellow, and purple, respectively), eventually draining into the vena cava (dark blue). Note that this color convention is used consistently throughout this article. The spatial extent of the liver is aligned horizontally with different liver segments tracked and labeled according to their one-dimensional (1D) spatial coordinates. We plot boxes around each liver segment with a width that is proportional to the user-specified volume fraction of that given segment. We also allow for user-specified fraction of outflow to the RHV, MHV, or LHV (as well as directly to the vena cava, as is the case for liver segment 1). Although variations of hepatic outflow have been described (21), for this study, we use standard liver anatomy for modeling and simulations. The standard parameters we use for liver volume fraction and outflow fraction are provided in Table 1.

In addition to the liver segment volumes and flow fractions, our model requires 21 parameters, from which we obtained 12 from literature, clinically measured eight from deceased donors (see next subsection), and implemented the remaining one as a free parameter; see Table 2. Unless otherwise specified, all simulations assume a mean pressure of 80 mmHg and 8 mmHg for the hepatic artery and portal vein, respectively, and an outlet pressure of 3 mmHg for the hepatic vein (where it meets the vena cava). We tested the sensitivity of the model to our choice of inlet/outlet pressure and verified a linear dependence for volume flow rate (e.g., a 5.0% increase in inlet pressures leads to a 5.0% increase in volume flow rates), indicating the model is fairly insensitive to this choice. The hepatic artery and portal vein continuously branch with a defined splitting number (i.e., the number of daughter vessels that emerge at each node) that is randomly sampled (and rounded) from a normal distribution with a user-specified mean and standard deviation; for the “ideal trifurcations” case presented below, a mean of 3 with standard deviation 0 is used, while for the “variable branching” case the splitting numbers presented in Table 2 are used. Any randomly generated branching values outside of the range 1 to 6 are discarded and, for each generation, the splitting number of the final branch is slightly increased or decreased to match the number of vessel segments in the next generation (our algorithm builds vascular trees from the microvasculature toward larger vessels).

**Table 2:** Parameters used in this study. All vessel radii reported in this study, except for the “Lumped sinusoid channel radius” are based on mean measurements in deceased liver donors; see Tables S1-S2.

| Parameter | Value | Reference |
| --- | --- | --- |
| Inlet pressure (hepatic artery) | 80 mmHg |  |
| Inlet pressure (portal vein) | 8 mmHg | (83) |
| Outlet pressure (hepatic vein) | 3 mmHg |  |
| Liver volume | 1518 cm<sup>3</sup> | (15) |
| Splitting number (hepatic artery) | 2.76±1.01 | (23, 36, 84) |
| Splitting number (portal vein) | 2.80±0.61 | (23, 36, 84) |
| Splitting number (hepatic vein) | 3.22±1.46 | (23, 36, 84) |
| Length ratio (hepatic artery) | 4.85 | (85) |
| Length ratio (portal vein) | 5.11 | (85) |
| Length ratio (hepatic vein) | 5 |  |
| Hepatic artery base lumen radius | 2.2 mm | This study |
| Hepatic artery terminal lumen radius | 3.0 μm | This study |
| Portal vein base lumen radius | 4.9 mm | This study |
| Portal vein terminal lumen radius | 25.1 μm | This study |
| RHV base lumen radius | 11.7 mm | This study |
| MHV base lumen radius | 4.0 mm | This study |
| LHV base lumen radius | 6.5 mm | This study |
| Hepatic vein terminal lumen radius | 28.8 μm | This study |
| Blood dynamic viscosity | 3.71 × 10<sup>-3</sup> Pa·s |  |
| Lumped sinusoid channel radius | 10.5 μm | This study |
| Lumped sinusoid channel length | 588 μm | (29) |

The terminal vessels of the hepatic artery and portal vein meet at the portal triads forming a confluence of flow directed through the sinusoids (green in Fig. 1C-D) which drains to a central hepatic vein. We model each sinusoid as an individual channel, 588 µm in length (29) connecting a portal triad to an adjacent central hepatic vein. Each centrilobular vein receives blood from six adjacent portal triads, and each portal triad feeds into three adjacent central veins (Fig. 1C), requiring that our model has twice as many terminal hepatic arteries and terminal portal veins as centrilobular veins. Although there are numerous parallel sinusoid channels connecting each portal triad to an adjacent centrilobular vein (30), we implement a “lumped sinusoid channel” which can be interpreted as an equivalent resistor that lumps together the net resistance of the local complex sinusoid network. This approach greatly increases the computational efficiency of our current model. Our choice of idealized connectivity captures an idealized, infinite hexagonal tiling of liver lobules, a segment of which is shown in Fig. 1D. The branches of the centrilobular veins merge with defined splitting numbers, eventually forming the base of the RHV, MHV, or LHV, which merge and drain into the vena cava; see color-coded schematic in Fig. 1B. We choose the number of generations *g* for each branching vascular tree (hepatic artery, portal vein, RHV, MHV, or LHV) based on the relation *s^g^* ≈ *N*, where *s* is the mean splitting number (Table 2) and *N* is the number of terminal vessels at the sinusoids. Since our parameters specify *s* and *N*, we compute *g* according to *g* = [log *N* / log *s*], where [·] denotes rounding to the nearest integer. The multiplicative change in radius γ between sequential generations is then chosen to satisfy the relation *r_f_* = γ^g^*r*_0_, where *r*_0_ is the base radius and *r_f_* is the terminal radius.

For both the inflow (hepatic artery/portal vein) and outflow (hepatic vein), we require that the two largest generations remain unchanged so that our model faithfully captures the most common vascular morphology, but for the third generation down to the sinusoids, we generate vascular networks either using idealized trifurcations or by randomly sampling Gaussian distributions of splitting numbers with the mean and standard deviation values indicated in Table 2 (randomly sampled values were rounded to integers). All liver models presented below contain a total of 5 million lobules in the unresected liver. This number is based on a reported total liver volume of 1518 cm^3^ (15), and an estimated volume of 0.3 mm^3^ per lobule, which we estimate based on the geometric mean of the limits of the lobule volume range of 0.1 to 0.9 mm^3^ reported by Teutsch (29).

We model the hydraulic conductance (inverse of hydraulic resistance) for every vascular segment connecting nodes *i* and *j* as laminar Poiseuille flow:

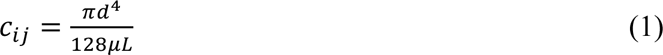

where *d* is the vessel diameter, *µ* is the dynamic viscosity, and *L* is the vessel segment length. We note that a resistive model based on Poiseuille flow is reasonable since inertial effects are negligible, based on the following estimates. For the largest vessel (base of the right hepatic vein), the Reynolds number is about 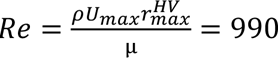 where ρ = 1050 kg/m^3, *U*_max_ = 0.30 m/s, 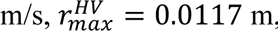, and µ = 3.71 × 10^67^ Pa·s (flow is laminar for *Re* ≲ 2000). For the most pulsatile vessel (base of the hepatic artery), the Womersley number is 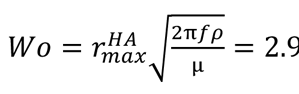, where 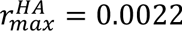 and *f* = 1 Hz is the human heart rate (for *Wo* ≲ 3, the velocity profile is parabolic and does not change shape over the cardiac cycle). We then solve the associated linear algebra problem *Cp* = *z* which is of the form:

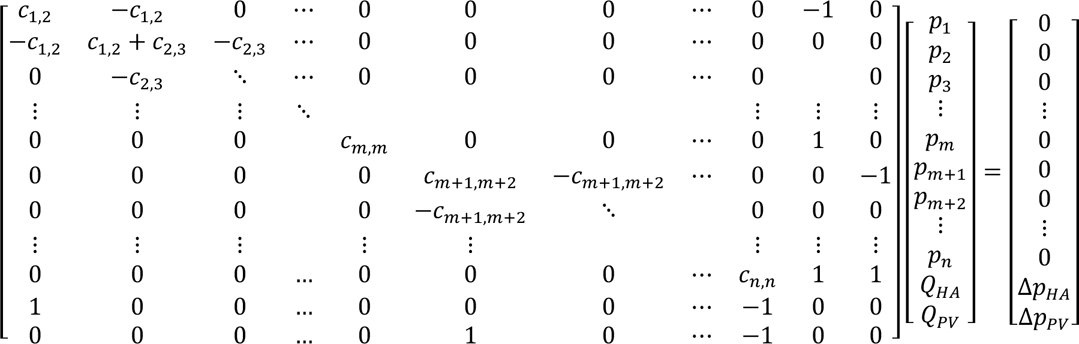

where *C* is a sparse matrix containing conductance values *c_i,j_*, *p_k_* are the unknown values of pressure at each node in the network, *Q_HA_* and *Q_PV_* are respectively the unknown total hepatic and portal volume flow rates, *m* is the total number of nodes forming the hepatic artery, *n* is the overall total number of nodes in the network, and *z* contains entirely zeros except for the last two entries Δ*p_HA_* and Δ*p_pv_* which respectively specify the hepatic and portal inlet pressures. We emphasize that for the large networks presented below, the total number of nodes is more than 2 × 10^7^, meaning *C* has over (2 × 10^7^)^2^ = 4 × 10^14=^ entries, so sparse storage of *C* is absolutely critical for computational tractability.

After obtaining the pressure, we compute the volume flow rate through every vessel segment as *Q_i,j_* = *c_,j_*(*p_i_* – **p*_j_*). We then computed the average blood flow speed *U* by dividing the local volume flow rate by the cross-sectional area of each given vessel. Finally, we computed the wall shear stress throughout the entire network as:

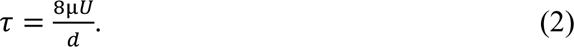

Both the construction of the spatial representation of the network and the linear algebra computations are performed using MATLAB (The Mathworks, Inc., Version R2021b); specifically, the vector *p* is solved using the backslash operator (i.e., *p* = *C*\*z*). Our largest reported networks model blood flow through ∼28 million nodes and 68 million individual vessel segments. Associated files are of order 1 GB and all associated calculations (network construction, solving for pressure, and computing volume flow rate, speed, and wall shear stress) are fast, requiring ∼12 min total on a standard laptop (Macbook Pro, 2019 13-inch with a 2.4 GHz Quad-Core Intel Core i5 processor).

For each virtual resection presented below (Fig. 3), we specify a polygon-shaped region of interest (ROIs; Fig. 3A-B) and obtain the matrix indices for all nodes inside of a given ROI. The corresponding nodes and all vessel segments contained partially or entirely within the ROI are then removed from the linear algebra system. Hence, there is no need to specify additional boundary conditions at the location of resection in our model. During resections comprising of segmental margins, any vessels that are large will be reconstructed to prevent significant outflow congestion. However, our modeling does not include reconstructions, but rather simulates physiological changes when there is complete venous outflow obstruction as a result of resection. In our resection simulations, we impose a fixed portal volume flow rate *Q_pv_* equivalent to the value of the unresected liver (and the fixed hepatic artery inlet pressure of 80 mmHg remains unchanged). The fixed portal volume flow rate boundary condition is achieved by solving a root-finding problem using the built-in MATLAB function “fzero.” We do so by varying the portal pressure in the range 4 to 100 mmHg to determine the value that satisfies 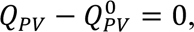, where *Q_pv_* and 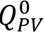 are the volume flow rates through the resected and unresected liver, respectively. Although this approach is computationally more expensive, requiring multiple iterations of solving the associated linear algebra problem, it has the benefit of simplicity in that it does not require an alternate formulation of the linear algebra problem.

Throughout this article, we plot the spatial representations of a given liver network with color indicating the volume flow rate, mean flow speed, or wall shear stress. Our 1D spatial representation of the liver facilitates interrogation of variations in hemodynamics across different liver segments, at the boundaries of adjacent segments, and across different branching generations, a feature which is utilized extensively below. We also plot the volume flow rate as a function of generation for each vessel. Since we model the sinusoids using the idealization of a single vessel of fixed length and diameter (specified below) rather than a complex network of parallel capillaries (30–33), we do not plot the mean volume flow rate for sinusoids. Our sinusoid modeling approach differs from several previous studies which have modeled sinusoidal flow as a porous medium (34–37); this difference is motivated further below. In the large networks presented below, we find that it is best to only plot the few generations corresponding to the greatest vessel diameters; plotting substantially more of the network is not computationally feasible nor are the details visually discernible. In such cases, we will replace several vessel generations with a white box labeling the liver segment (e.g., Fig. 2A).

**Fig. 2:**
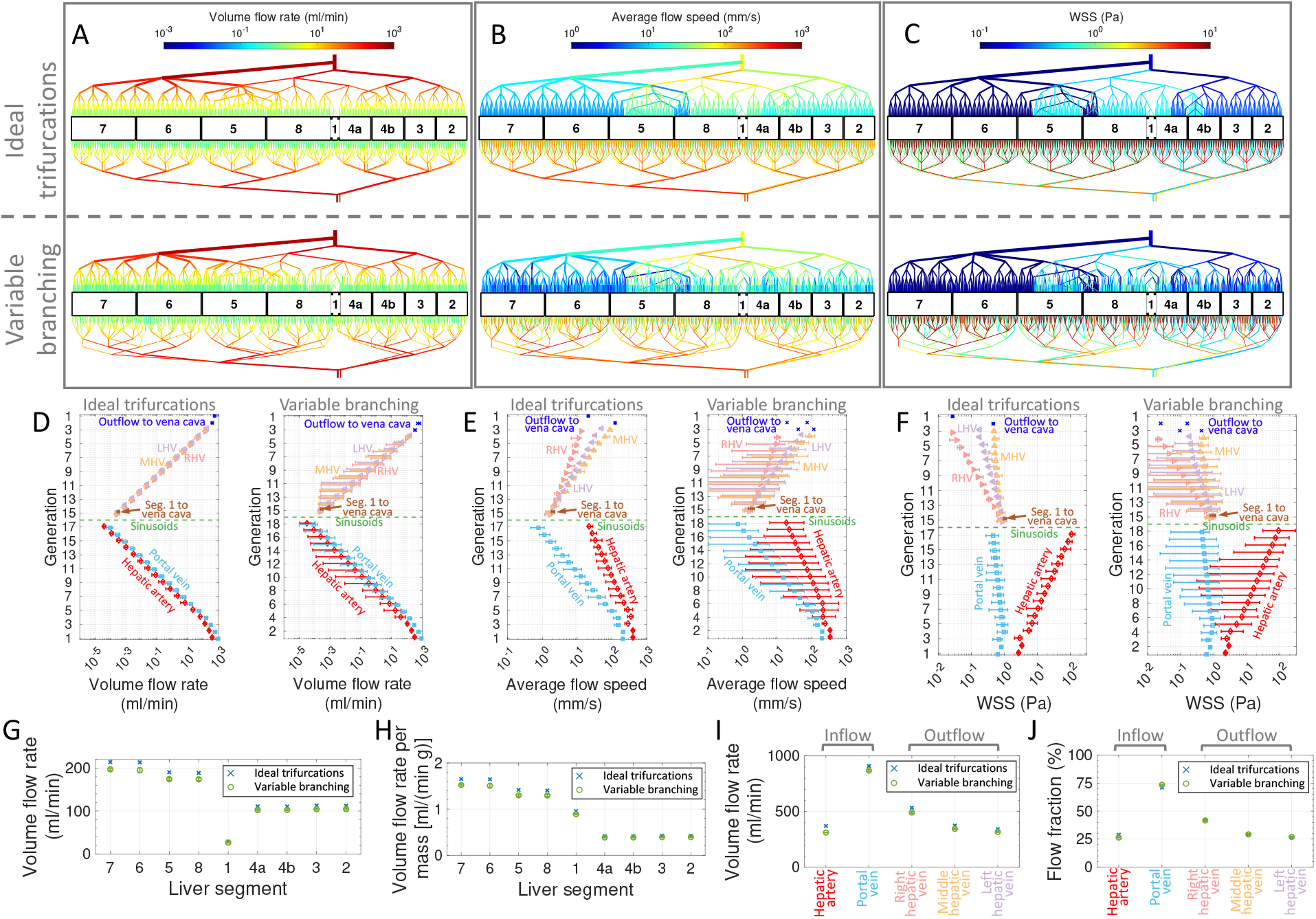
Comparisons of idealized trifurcations or realistic variable branching networks. (A-C) Schematics of realistically-sized networks (each containing 5 million lobules) showing only the first few branching generations, with color encoding the volume flow rate, average flow speed, and wall shear stress (WSS), respectively. The top and bottom rows correspond to idealized trifurcations and realistic variable branching, as indicated. (D-F) Semilogarithmic plots of the branching generation versus the volume flow rate, average flow speed, and wall shear stress with different vessel segments color-coded and labeled. The left and right plots correspond to idealized trifurcations and realistic variable branching, as indicated. Quantities are not plotted for the lumped sinusoid channels (labeled “sinusoids”), which are idealized (see Methods). The uncertainty bars indicate the range of the data. (G-J) Plots of the volume flow rate and volume flow rate per unit mass (i.e., hepatic artery and portal vein) for (G-H) each of the nine liver segments and (I-J) each of the five major inflow/outflow vascular networks, as indicated. The uncertainty bars for the variable branching correspond to the standard deviation across 10 simulations, indicating results are robust for the randomly generated variable branching networks. The volume flow rates and flow fractions closely approximate clinical observations.

### Clinical measurements of deceased donor livers

Measurements were obtained from 12 healthy livers of deceased donors that were used for transplantation (Tables S1-S2). The diameter of the main trunk of the portal vein was measured using a sterile intra-operative measuring ruler 3 mm proximal to the bifurcation of left and right portal veins. The left and right portal veins were measured 3 mm distal to the bifurcation. Similarly, the diameter of the hepatic artery and its two branches were measured using the same protocol as that of the portal vein. Liver biopsies obtained from respective donors (as per clinical routine protocol to assess hepatic steatosis) were used by a liver histopathologist to measure the terminal branch diameter of the portal vein, hepatic artery, central vein, and vessel wall thickness. The lumen radius for each given vessel (Table 2) was then calculated as *r* = (*D* − 2*w*)/2, where *D* is the measured diameter of a given vessel (Table S1) and *w* is the vessel wall thickness (Table S2).

## Results

### Variation in splitting number alone generates substantial flow heterogeneity

Prior studies have often assumed that the hepatic artery, portal vein, and hepatic vein branch as perfectly ideal trifurcations (i.e., with splitting number 3) (36). Thus, we first investigated the effect of implementing realistic, randomly sampled splitting numbers in our simulations. Visualizations of the volume flow rate, average flow speed, and wall shear stress for the ideal trifurcations and variable branching cases are provided in Fig. 2A-C, which immediately reveal greater heterogeneity in the variable branching case. We found that the total number of nodes and vessel segments was nearly identical for the two different approaches across a wide range of total number of lobules in the network, and importantly, we also found that the additional calculations required for implementing variable branching increased the computational time minimally (Supplemental Fig. S1A) for a fixed number of lobules. We furthermore confirmed that our randomly generated branching networks did indeed have a mean and standard deviation for the splitting number that closely matches the intended splitting numbers available in Table 2 (Supplemental Fig. S1B).

To quantify the differences in heterogeneity in the two cases, we plotted the mean and range of each quantity (volume flow rate, mean speed, and wall shear stress) as a function of generation in Fig. 2D-F. These plots reveal that mean quantities for the three cases are remarkably similar, with a couple small differences worth highlighting. First, the number of generations for the ideal trifurcations case is one fewer for the hepatic artery and portal vein, but one greater for the hepatic vein (Supplemental Fig. S1C); this is a consequence of using a splitting number of 3 in the ideal case, which is respectively greater and lower than the value used in the variable branching case (see Table 2). Secondly, the ideal trifurcation case is not perfectly homogenous across parallel vessels and instead does exhibit a relatively small amount of variability at a given generation (i.e., the standard deviation is nonzero as indicated by the uncertainty bars). This has two causes: (i) LHV, MHV, and RHV have different diameters based on our measurements (Table 2), and (ii) we assume a heterogeneous branching geometry for the first two generations (i.e., those with largest radius) of the hepatic artery, portal vein, and hepatic vein. To test whether the volume flow rates really are quite similar between simulations with ideal trifurcations and those with variable branching – and if there is substantial variation across different realizations of the randomly generated variable branching case – we compute the total volume flow rate for each liver segment in each case (Fig. 2G). We found that the volume flow rate for each liver segment in the ideal trifurcation case was only 6 to 9% larger than that of the variable branching case on average, and the mean volume flow rates across different variable branching cases were negligibly small (the uncertainty bars in Fig. 2G are smaller than the size of the symbols). These simulations indicate that variability in the splitting number of the network greatly increases the heterogeneity of the flow across parallel branches, with minimal impact on the overall mean volume flow rate through the entire liver. This increased heterogeneity can be understood purely as a result of decreased hydraulic resistance arising for parallel flow routes with larger numbers of branches; we illustrated this concept with a simple analytical example in Fig. S2.

### Simulations predict realistic perfusion rates and flow distribution

Our simulation is based entirely on parameters either reported in the literature or that we have clinically measured, except for one parameter: the radius of the lumped sinusoid channels (each of which we model as a single vessel). We used this quantity as a free parameter to tune the total volume flow rate, and we find that for a radius of 10.5 µm, we obtained overall volume flow rates of 1280 ml/min and 1180 ± 10 ml/min for the cases of ideal trifurcations and variable branching, respectively. We also computed the overall volume flow rate per unit mass (i.e., hepatic artery and portal vein), which is plotted in Fig. 2H and ranges from about 0.5 to 1.5 ml/(min·g) which agrees well with reports of overall mean liver perfusion per unit mass of about 1 ml/(min·g) in the literature (38, 39). Finally, we analyzed distributions of inflow and outflow among the hepatic artery/portal vein and the RHV/MHV/LHV, which are plotted in Fig. 2I. We converted this to a plot of the overall flow fraction, which is shown in Fig. 2J and confirms that our model predicts realistic flow distributions with about 25% of perfusion inflow provided by the hepatic artery and 75% provided by the portal vein (38). The dominant source of outflow is RHV, followed by the MHV, then the LHV, in agreement with clinical characterization of the most prevalent liver anatomy (40).

### Quantification of hyper-perfusion for different liver resections

Having confirmed that our model makes quite accurate predictions of perfusion for an intact liver, we next investigated alterations in hemodynamics resulting from different clinically performed liver resections. We considered one scenario of left hepatectomy and three scenarios of right hepatectomy (Fig. 3A-F). In the former case, blood outflow will distribute throughout the entire LHV and the remaining portion of the MHV. However, we consider three different scenarios in the latter case: two in which the entire left liver is retained and the main trunk of the MHV is either retained (Fig. 3D) or resected (Fig. 3E), or a scenario of extended right hepatectomy in which only liver segments 2 and 3 remain, with outflow routing through the remaining portion of the LHV (Fig. 3F). We numerically performed these resections by removing the associated nodes and vessel segments from the network for each given scenario (so no additional boundary conditions are specified), then recomputed the volume flow rate under the assumptions of a fixed hepatic artery inlet pressure of 80 mmHg and a fixed portal volume flow rate *Q*_;5_equivalent to the value of the unresected liver (see Methods). The latter boundary condition is justified because the entirety of blood from the bowel and spleen is directed through the superior mesenteric vein and the splenic vein respectively, forming the portal vein; there is minimal variation in this volume flow rate, except under pathological rerouting of blood or due to the effect of anesthesia intra-operatively affecting the flow from the splanchnic system, both of which are beyond the scope of this paper. We highlight that our boundary condition assumptions lead to portal hypertension, which is clinically well-established (41) and discussed in detail further below.

**Fig. 3:**
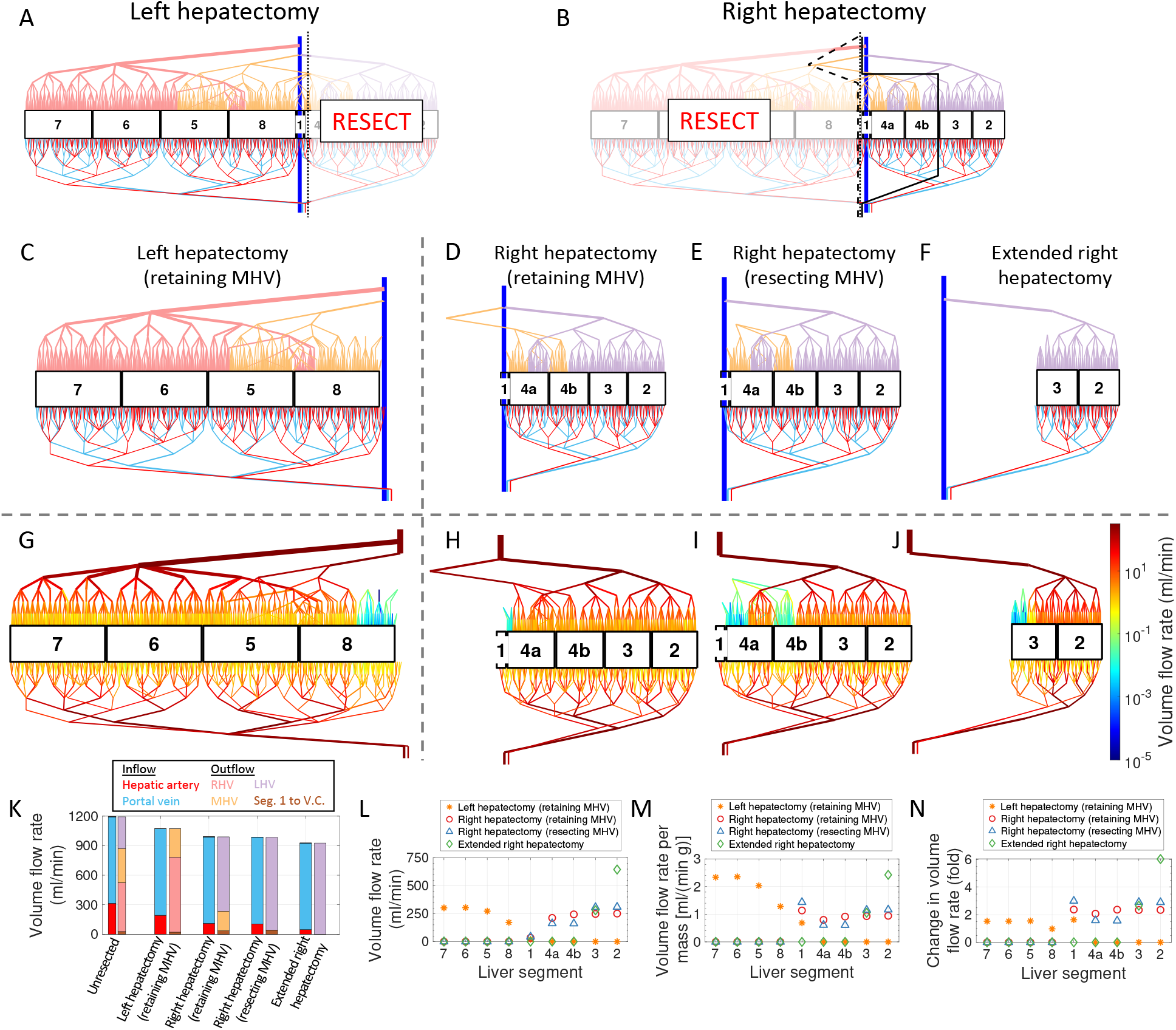
Simulations of variable liver resections predicting hepatic hemodynamic alterations. (A-B) Schematics of left and right hepatectomy applied to a realistically-sized network (containing 5 million lobules); dashed/dotted/solid black lines indicate surgical resection planes. (C-F) Schematic illustration of networks following (C) left hepatectomy (33% resection) or (D-F) three variations of a right hepatectomy (67%, 67%, and 84% resection, respectively), as indicated. (G-J) Schematic plots with color, encoding the volume flow rate. The color bar at the far right corresponds to all four plots. (K-N) Plots of the (K-L) volume flow rate, (M) volume flow rate per unit mass, and (N) fractional change in volume flow rate for different (K) vessels and (L-N) liver segments under all four resection scenarios.

To quantify changes in liver perfusion following resection, we plotted the total volume flow rate for the inflow and outflow in each case (unresected and each of the four resections) as shown in Fig. 3K. This plot indicates that increasingly larger resections (i.e., smaller remaining portions of the liver) lead to reduced total volume flow rate. Because we assume the portal inflow *Q_pv_* is fixed, this inflow reduction is entirely attributable to decreased hepatic inflow (red bar in Fig. 3K) arising from increasing resistance of the liver as greater portions are removed. The distribution of outflow follows logically from the portions of the hepatic vein that remain. For example, left lobe outflow via the MHV is preserved for segments 4a/b after a right hepatectomy in which the MHV is not resected (Fig. 3H). However, those portions of segment 4 primarily draining through the MHV become congested with MHV resection (Fig. 3I). In Fig. 3L-M, we plot the volume flow rate and volume flow rate per unit mass for all four resection scenarios. These plots, compared to those of the unresected liver (Fig. 2G-H), indicate that perfusion is elevated above the baseline value for at least one liver segment in all resection scenarios. To quantify the change in volume flow rate, we plotted the ratio of the volume flow rate for each liver segment in a given resection scenario to that of the unresected liver in Fig. 3N. This plot reveals a typical two-to three-fold perfusion increase in the remaining liver segments, but this hyper-perfusion may be as high as a six-fold increase in the case of the extended right hepatectomy. To test the consistency of our observations across different randomly generated variable branching liver models, we performed analogous liver resections for several more networks and confirmed the robustness of our observations; Fig. S3 shows a replication of Fig. 3 for a second random network. The only considerable change we observe is moderate variability in the perfusion of individual liver segments; we next describe the origin of this variability, as it relates to outflow obstruction.

Obstructed outflow induces complex hemodynamic alterations. Compared to the unresected network (Fig. 2A bottom), every resection contains a region of the hepatic vein with greatly reduced volume flow rate (blue/green portions of Fig. 3G-J). The cases of right hepatectomy in which the MHV is either retained or resected (compare Fig. 3H and I) clearly indicate such regions of reduced flow arise from obstructed outflow due to resection: the former case shows a small region of reduced flow mostly localized to the portion of segment 4a adjacent to segment 1, while the latter case exhibits a much greater obstructed region in the left portions of both segments 4a and 4b (these portions drain through the MHV in the former case). To investigate alterations in hemodynamics arising from obstructed outflow, we plotted the ratios of the volume flow rate of the right hepatectomy for scenarios of both retained and resected MHV, to those of the unresected liver in Fig. 4A-B. These plots reveal many important hemodynamic changes, which we will describe sequentially starting with the outflow, then the sinusoids, and finally the inflow.

**Fig. 4:**
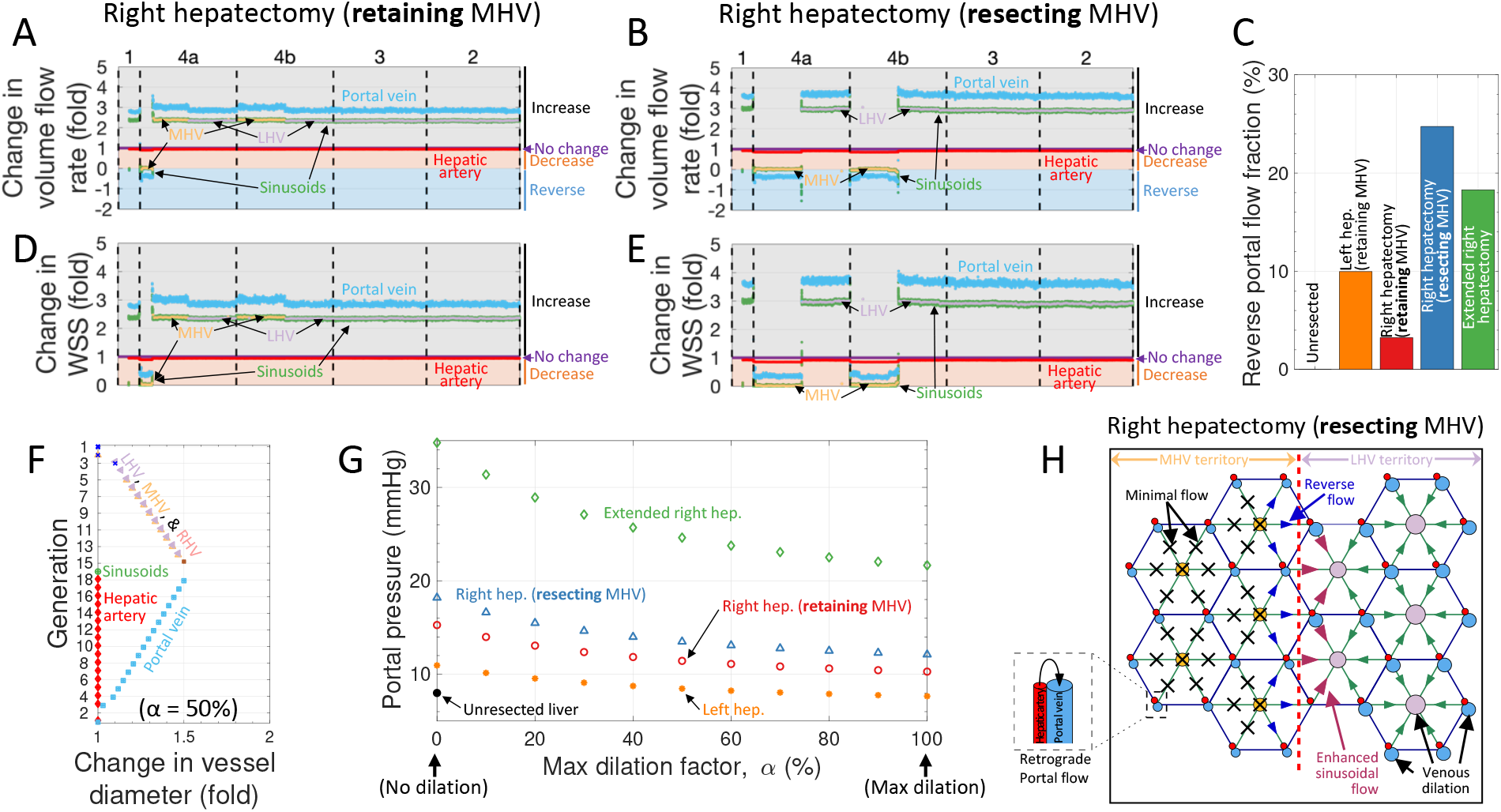
Quantification of perfusion alterations for variable liver resections. (A-B) Plots of the factor by which volume flow rate changes in the resected liver, compared to the unresected liver, for each vessel (color-coded) in each liver segment (labeled and divided by dashed black lines) following right hepatectomy with MHV (A) retention or (B) resection. Labels at the right indicate regions in which the resected liver volume flow rate increases, is unchanged, decreases, or reverses. Quantities are plotted seven generations from the sinusoids. (C) The total percentage of portal vein segments which exhibit retrograde flow in each indicated resection scenario. (D-E) Plots analogous to (A-B), but for wall shear stress. (F) A plot of the dilation, for α = 50%, as a function of generation applied to each color-coded vessel. Note that dilation is implemented for only the portal and hepatic veins. (G) Plots of the portal pressure as a function of the maximum dilation factor α for all four resections. (H) A schematic illustrating the key features of altered hemodynamics obtained in our model for right hepatectomy with MHV resection. Note that the corresponding unresected liver has 5 million lobules, and lumped sinusoid channels are labeled only as “sinusoids” for the sake of concise labeling.

Unsurprisingly, we find that the case of right hepatectomy with MHV retention exhibits an approximately uniform ∼2.5-fold elevated volume flow rate throughout both the MHV and LHV (Fig. 4A); this elevation is primarily a consequence of perfusing the same portal blood supply through a smaller residual liver. However, in the case of MHV resection, there is zero volume flow rate through the MHV and flow through the LHV is even greater than in the former case (elevated ∼3-fold; Fig. 4B). This further elevated flow through the LHV in the latter case arises due to the substantial obstruction of outflow that occurs when the MHV is resected, routing even more blood through the LHV.

Flow through the lumped sinusoid channels exhibits largely the same phenomenology as the outflow through the MHV and LHV for both cases, which is a consequence of continuity (i.e., flow cannot enter the sinusoids if it cannot continue through to the resected MHV). However, for the case of MHV resection, the lumped sinusoid channels exhibit “spikes” in volume flow rate near the boundary of the territory in which blood either drains to the LHV or MHV. These spikes are positive (indicating anterograde flow) on the LHV-side of the boundary and negative (indicating retrograde flow) on the MHV-side of the boundary. This indicates that at the boundary of the vascular territory for (obstructed) MHV outflow and (unobstructed) LHV outflow, there is substantial reversal of blood flow through the sinusoids which then drains anterograde though neighboring sinusoids, further elevating their perfusion by as much as ∼50% of the baseline value. Of all the vessels, the hepatic artery exhibits the least variation in local volume flow rate, remaining nearly the same as the unresected liver in all cases, except for a small reduction in any territory with obstructed outflow (e.g., half of 4a and 4b in Fig. 4B). Conversely, the left portal vein exhibits the greatest increase in volume flow rate of all vessels, which is about 3-fold and 3.7-fold after a right hepatectomy with retained or resected MHV, respectively. Interestingly, when the MHV is resected obstructing outflow, the portal vein exhibits low magnitude retrograde flow. Since the flow through the sinusoids is zero through most of this region, this retrograde flow originates from the hepatic artery. We confirmed that these same trends persist throughout most branching generations, albeit with increasing variability for the portal vein flow across sequential generations (Fig. S4). We furthermore quantified the percentage of all portal vein segments (across the entire network) that exhibit retrograde flow, which is plotted in Fig. 4C for all four resections and the unresected case. All resections exhibit some portal retrograde flow, but the left hepatectomy and extended right hepatectomy have intermediate values of 10% and 18%, whereas the right hepatectomy without MHV resection has a mere 3% retrograde flow percentage, while resection of the MHV leads to a substantial 25%. Hence, resection with obstructed outflow (e.g., MHV resection) not only increases perfusion through the patent route (e.g., LHV), but also generates hyper-perfusion and retrograde flow through the sinusoids at the boundary of the territories, with unobstructed and obstructed outflow, respectively, as well as retrograde portal flow sourced from the hepatic artery in the territory of obstructed outflow.

The ratio of wall shear stress in two resected livers to that of the unresected liver is plotted for different vessels across different liver segments in Fig. 4D-E. Since wall shear stress is linearly proportional to the volume flow rate (equation (2)), these plots exhibit many features that qualitatively match that of the volume flow rate, such as dramatic, nearly uniform increases for the MHV, LHV, and portal vein for the right hepatectomy with MHV retention. In the case of MHV resection, wall shear stress in the LHV and portal vein is further elevated, again due to the obstructed MHV outflow.

### Acute venous dilation reduces portal hypertension

Prior studies (17, 42) have noted that following liver resection, venous dilation may occur to compensate for elevated hydraulic resistance arising from the reduction in liver volume (i.e., liver resection reduces the number of parallel vessels). To test this hypothesis, we implemented a scheme for a gradual increase in venous dilation wherein the extent of dilation varies linearly with generation, with no dilation of the largest veins and a maximum dilation percentage α for the smallest veins, where 0%≤ α ≤ 100%. Note that we impose zero dilation for the entire hepatic artery and lumped sinusoid channels. The extent of dilation as a function of generation, in the case of α = 50%, is shown in Fig. 4F. Importantly, hydraulic resistance is strongly dependent on vessel diameter (proportional to d^-4^), so we next investigated whether venous dilation would reduce portal pressure. Fig. 4G shows the portal pressure as a function of α for the four different resections. In all cases, greater venous dilation leads to a monotonic decrease in portal pressure, but the effect is more dramatic for greater resection volumes. The extended right hepatectomy, in particular, exhibits a portal pressure of 34.8 mmHg with no dilation, but that pressure reduces to 24.6 mmHg when α = 50% and 21.7 mmHg when α = 100%. Comparing the two cases of right hepatectomy reveals that without venous dilation, portal pressure rises to 15.3 mmHg if the MHV is retained, whereas the case with MHV resection further elevates the pressure to 18.2 mmHg. The maximum venous dilation we test (α = 100%) leads to portal pressures of 10.3 and 12.1 mmHg for these same respective cases. Finally, the left hepatectomy elevates portal pressure to 11.0 mmHg with no venous dilation, and maximum dilation (α = 100%) leads to a portal pressure of 7.7 mmHg, similar to the assumed baseline value of 8 mmHg for the unresected liver.

## Discussion

The liver has significant, but limited ability to regenerate after resection. It has been taught that between 70-80% of the normal liver can be safely resected (43, 44) with confidence that the liver will regenerate. However, the mechanisms and science behind this limit of “safe” resection are unclear. Liver perfusion pressures have been correlated with liver regeneration and increased portal pressures have been associated with liver injury. This study introduces a comprehensive model of blood flow through the entire human liver which makes initial strides toward uncovering limits of safe resection. Our approach utilizes lumped parameter modeling which has enabled computation of pressures, volume flow rates, mean flow speeds, and wall shear stresses throughout an entire network of 5 million lobules, consisting of over 28 million nodes and over 68 million vessel segments. These calculations run in just over 12 minutes on a standard laptop. Our model is built on realistic anatomy-based parameters available in the literature, as well as vessel radii which we have clinically measured in livers of deceased donors. The most novel aspects of our model include the demarcation of liver segments and the incorporation of realistic vessel branching morphology. In the latter case, we explicitly specify the connectivity of the two largest generations of portal vein, hepatic artery, and hepatic vein, then generate dendritic trees with sequential branching numbers sampled from clinically measured statistical distributions to connect these macroscopic vessels to the microscopic sinusoids. Our model for the intact liver predicts volume flow rates in good agreement with clinical measurements (Fig. 2G-J).

Having demonstrated the accuracy of predictions from our intact liver model, we next simulated variable liver resections (Fig. 3C-J) and quantified the change in volume flow rate (Fig. 3K-N). We emphasize that this model is intended to capture acute hemodynamic alterations and does not capture the gradual restoration of normal hemodynamics that result from regeneration of the remnant liver. Since liver resection removes a fraction of parallel vasculature proportional to the resected mass fraction, the hydraulic resistance of the liver correspondingly increases. This has two distinct effects on the hepatic artery and portal vein, for which we assume constant inlet pressure and constant inlet volume flow rate, respectively. In the former case, increasingly severe arterial hypoperfusion occurs with greater mass resections, a prediction which is corroborated by experimental measurements (45). In the latter case, the increased resistance with fixed portal inflow results in portal hypertension, which is more severe with greater mass resection (i.e., greater resistance; Fig. 4I). Prior experimental studies also confirm correlation between the extent of mass resection and portal hypertension severity (46). One additional consequence of a constant portal inflow is that, despite the arterial hypoperfusion, there is an overall increase in the post-resection volume flow rate in remaining liver segments and the volume flow rate per unit mass (Fig. 3K-N). Our model predicts perfusion increases that are often two-or three-fold, and as high as six-fold for the extended right hepatectomy (segment 2 in Fig. 3N). These predictions are supported by prior direct flow measurements following partial hepatectomies which have recorded several-fold increases in the portal flow per tissue volume unit (47). In particular, extended hepatic resection remains a high-risk procedure with potential postoperative hepatic dysfunction and progressive liver failure imposing high morbidity and mortality (48). We highlight that the cases of right hepatectomy (either retaining or resecting the MHV) involve approximately the same anatomic mass removal; however, the latter scenario exhibits much greater prevalence of stagnant flow in the hepatic veins (Fig. 4A-B). Consequently, the latter case leads to a substantially greater increase in perfusion of sinusoids draining to the LHV (approximately three-fold; Fig. 4B) compared to the former case (approximately 2.4-fold; Fig. 4A). It is also notable that outflow obstruction may lead to local retrograde rerouting of arterial blood through the portal vein (Fig. 4B). Consequently, the effected liver territory would likely exhibit diminished functional capacity due to the diminished sinusoidal flow and lack of nutrient-rich portal blood. Indeed, for the left hepatectomy, right hepatectomy (with MHV retention), right hepatectomy (with MHV resection), and extended right hepatectomy, the liver volume is respectively reduced to 66.6%, 33.4%, 33.4%, and 15.9% of its original volume, but only 61.3%, 33.2%, 25.1%, and 15.6% of the volume has substantial flow through the sinusoids. Hence, our model supports the notion of a “functional” liver volume which may be reduced beyond that of the anatomic mass due to flow disruption in some regions (49, 50).

In addition to volume flow rate and average flow speed, our simulations also provide quantitative predictions of elevated wall shear stress. Elevated shear stress has long been hypothesized to stimulate liver regeneration after partial resection (51, 52) or liver transplant with a size mismatch between recipient and donor (53). Indeed, as early as one minute after partial hepatectomy in rats, Mars et al. observed the appearance of urokinase receptors, which activate hepatocyte growth factor (HGF), initiating liver regeneration (54). More recently, Ishikawa et al. provided direct evidence that changes in sinusoidal wall shear stress and tension mediate HGF production, signaling the initiation and termination of liver regeneration (30). Restoration of hepatic volume after partial resection is critical for maintaining metabolic function, and such regeneration involves proliferation of parenchymal and non-parenchymal cells throughout the liver (55–57). Our simulations of right hepatectomy with MHV retention (Fig. 4D) predict that the extent of wall shear stress increase is approximately uniform throughout all remaining sinusoids of the liver; this result suggests that after partial hepatectomy, the entire remaining liver participates in producing molecules that signal liver regeneration. On the other hand, right hepatectomy with MHV resection (Fig. 4E) exhibits near-zero wall shear stress for portions of liver segments 4a and 4b, in the vicinity of the MHV. Liver resections with MHV resection have been well documented to result in focal hepatic venous outflow obstruction that leads to formation of necrosis in the undrained tissue worsening the remnant liver mass (58). These insights highlight the importance of retaining the MHV in right hepatectomy.

This model provides insights into the acute hemodynamic alterations following various liver resections. As mentioned above, portal hypertension results as a direct consequence of a constant portal flow rate perfusing a smaller (more resistive) resected liver. Prior experimental studies have observed that this immediate increase in portal pressure following liver resection is directly correlated with (and therefore indicative of) elevated sinusoidal shear stress (41, 59). Clinically, increased intra-operative portal pressure beyond the threshold of 22 mmHg after partial hepatectomy is associated with liver failure and mortality (60). Our simulations predict that the extent of portal hypertension is below this approximate threshold for all resections considered, except the extended right hepatectomy (Fig. 4G). Our incorporation of venous dilation (Fig. 4F-G), which has been documented following embolization (61) quantifies a mechanism by which the liver may acutely reduce portal hypertension. For the functional form and extent of dilation assumed here, we predict that venous dilation may reduce portal pressure by as much as 38% (extended right hepatectomy in Fig. 4G). Surgical techniques to reduce portal hypertension involve ligating splenic artery and/or splenectomy (62), but this additional procedure often leads to complications (63–65). We hypothesize that these procedures can be avoided through pre- and intra-operative modeling to predict hemodynamic alterations associated with hepatic resection.

The liver has the most complicated hemodynamic circulation of an any organ. Blood flow through the liver is controlled by mechanisms independent of extrinsic innervation or vasoactive agents regulating hepatic artery and portal venous inflow. However, there is also an interrelationship between the two circuits with the hepatic arterial vasculature exhibiting autoregulation of blood flow. The autoregulation is used to describe the non-linearity of the arterial pressure-to-arterial flow relationship and is specific to local blood flow to remain constant when the pressure changes within the organ (66–70). The degree of autoregulation is considered small (70) and present in about 60% of the all liver resections (68). The observed effects are primarily mediated by myogenic adaptations of the vasculature to changes in transmural pressure (70). A second form of intrinsic regulation termed hepatic artery buffer response (HABR) is a unique mechanism representing the ability of the hepatic artery to produce compensatory flow changes in response to changes in the portal venous flow. With reduction in portal flow, the hepatic artery dilates and vice versa (71, 72). The physiological role of HABR is to minimize the influence of portal venous flow changes on the hepatic clearance and maintain adequate oxygen supply to tissues.

The increased blood flow to the liver soon after partial hepatectomy and the resultant increased intrahepatic shear stress have been proposed to stimulate and regulate liver regeneration (51, 52, 73). Following right lobe resections, the grafts are subjected to more than twice the portal flow. In the absence of active regulation, arterial flow might be expected to double as well, however, there is a striking decrease in arterial flow in right lobe grafts (74). Although HABR plays an important role in maintaining the blood flow to the liver, it is beyond the scope of this simulation. Capturing the effect requires inclusion of adenosine and nitric oxide via ATP-dependent stimulation of endothelial purinergic receptors in the hepatic artery. In the future, we hope to further incorporate HABR and develop our model through intra-operative measurements of arterial to portal venous flow and pressure ratios following resections.

There are numerous noteworthy limitations and opportunities for future improvement of our model. Perhaps most prominently, we have not yet validated our simulation against clinical measurements. A further limitation is that we have approximated the liver anatomy as a one-dimensional array of adjacent liver segments. Although this approach does improve the clarity and simplicity of data visualization and virtual liver resection, the tradeoff is a simplification of vessel connectivity to different liver segments. For example, liver segments 4b and 5 are adjacent (Fig. 1A) meaning branching microvasculature may connect to both segments, which is not captured in our model. Furthermore, our model does not capture the true tortuous and overlapping vascular geometry in 3D which renders clinical liver resection substantially more complex. A more realistic 3D geometry could perhaps be achieved by leveraging algorithms for generating synthetic vascular trees (75). A further simplification is that we model each local sinusoid network as a single channel with a 10.5 μm radius (a free parameter used to tune the overall liver resistance such that a realistic overall volume flow rate is obtained). Future models could implement more realistic sinusoidal anatomy, perhaps based on novel imaging approaches (30–33), which would enable reliable estimates of the volume flow rate and wall shear stress in individual sinusoidal vessels. The model would also benefit from prescribing the distribution of flow from the eight Couinaud segments to the right, middle, and left hepatic vein (Table 1) based on clinical measurements. Future simulations could also incorporate angiogenesis and oxygen extraction in a hyper-perfused remnant to predict the evolution of liver regeneration. We anticipate that inclusion of network heterogeneities (e.g., variable branching, as included here) would be important for the accuracy of such a model. All quantities associated with our model should be interpreted as time-averaged (e.g., mean arterial pressure, mean wall shear stress), but future simulations could implement a systolic/diastolic variation, as in prior studies (24, 26), to estimate changes in perfusion and peak wall shear stress across the cardiac cycle. Yet another limitation is that we have neglected any acute arterial response that may occur following liver resection (45), as well as the effects of anesthesia and any drugs that may be administered during liver resection that result in portosystemic shunting and splanchnic dilatation altering the portal flow being measured intra-operatively. Hepatic resections and surgical manipulation can cause reflex vasoconstriction which needs to be considered as a factor that can alter measurements. Finally, our pressure calculations assume Poiseuille flow and do not account for the non-Newtonian nature of blood, nor do they capture any minor resistive losses. Future simulations could incorporate more realistic scaling laws for resistance through microvasculature (based on an apparent blood viscosity as a function of lumen diameter (76)) and perhaps utilize results of 3D direct numerical simulations and/or 4D magnetic resonance imaging (MRI) to improve blood flow modeling in the largest vessels (77, 78).

Overall, this simulation constitutes an important incremental stride towards a quantitative patient-specific model that can help support pre- and intra-operative decisions and outcomes. Current preoperative diagnostics allow a comprehensive assessment of residual liver anatomy and function that includes liver volume, liver segment volume fraction, and large vessel anatomy to every segment, which can all be implemented in our model in a straightforward manner. We may further include velocity measurements in primary vessels (hepatic artery, portal vein, and hepatic veins) obtained via 4D flow MRI (79) to validate model predictions in the pre-operative planning for living donor liver transplantation or extensive resections. Implementation of patient-specific imaging modalities into the liver modeling during pre-operative surgical planning is critical for translational work. The current modeling is suited for living donor liver transplantation where the liver’s microscopic and macroscopic architecture is physiologically and histologically normal. For oncologic resections, further development of the model to incorporate biopsy directed liver simulations would be essential and is currently beyond the scope of this study. In cases of extended resections, portal flow is also assessed apart from metabolic functions of the remnant liver, to confirm the safety of the extent of resection. Through high resolution imaging, measurements of hepatic hemodynamics have also played a limited role in pre-operative liver assessments (80). Any number of measurements of vessel radii are straightforward to implement in our model, and volume flow rate / flow speed measurements can be implemented as constraints that improve the accuracy of our model. Ultimately, the most valuable aspect of numerically modeling liver resection is the ability to predict post-surgical alterations to perfusion. Such alterations impose traction, tension, and shear forces on liver tissue that play a critical role in regeneration. However, in clinical settings, such alterations cannot be determined or quantified pre-operatively. In future studies, we plan to validate predictions of our model against post-operative clinical measurements. Water-fat quantification techniques using MRI are available to measure tissue perfusion with quantitative characterization of parenchymal and microcirculatory changes (81) and to study the extent of liver damage (81, 82).

## Authors’ contributions

JT developed the mathematical liver model, performed all simulations, and wrote the manuscript; TLP overlooked the project and provided expert opinion; JSR conceived of the idea, obtained liver vascular measurements, and wrote the manuscript.

## Code and data availability

The simulation code developed for this study is publicly available online at https://doi.org/10.5281/zenodo.7490383. The specific data used to generate the plots in this manuscript are available from the authors upon request.

---

**Fig. S1:**
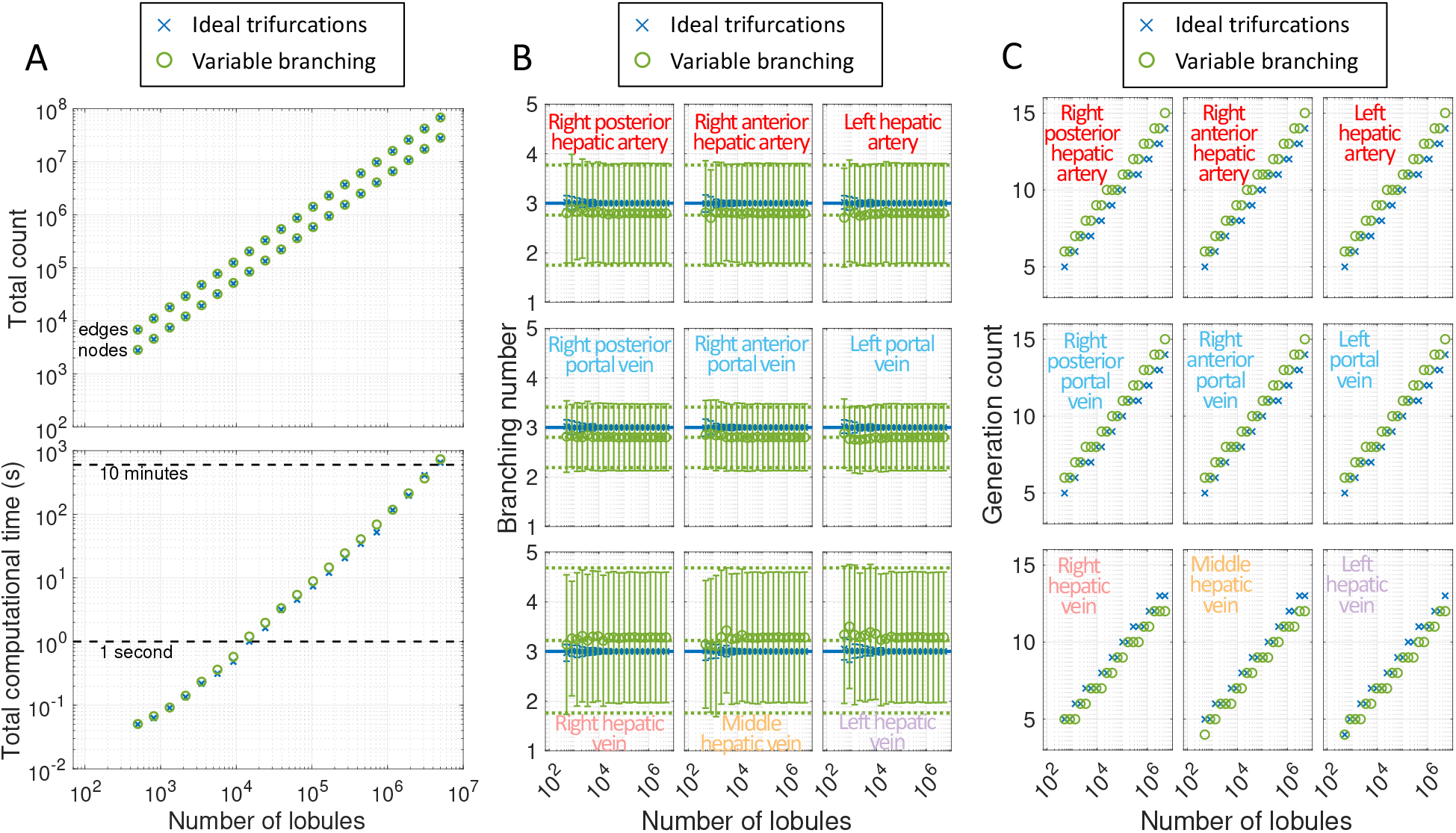
Scaling of various quantities as a function of number of lobules. (A) Plots of the total number of edges (i.e., vessel segments) and nodes (top plot) and the total computational time (bottom plot) as a function of the total number of lobules simulated. The differences between the two cases are marginal. (B) Plots of the splitting number as the total number of lobules in the network increases. The solid blue line indicates an idealized branching number of 3, and the dotted green lines indicate the mean and standard deviation for the variable branching, as indicated in Table 2. The circle markers and uncertainty bars indicate the mean and standard deviation from our network models, respectively. For small networks with idealized trifurcations, a substantial fraction of the branches may not be idealized trifurcations if the total number of lobules is not an integer power of 3, which explains why the blue uncertainty bars are a bit larger at low numbers of lobules. For the variable branching case, the actual mean and standard deviation is quite close to the values specified in Table 2. (C) Plots of the total number of vessel generations for a given vascular network as a function of the total number of lobules.

**Fig. S2:**
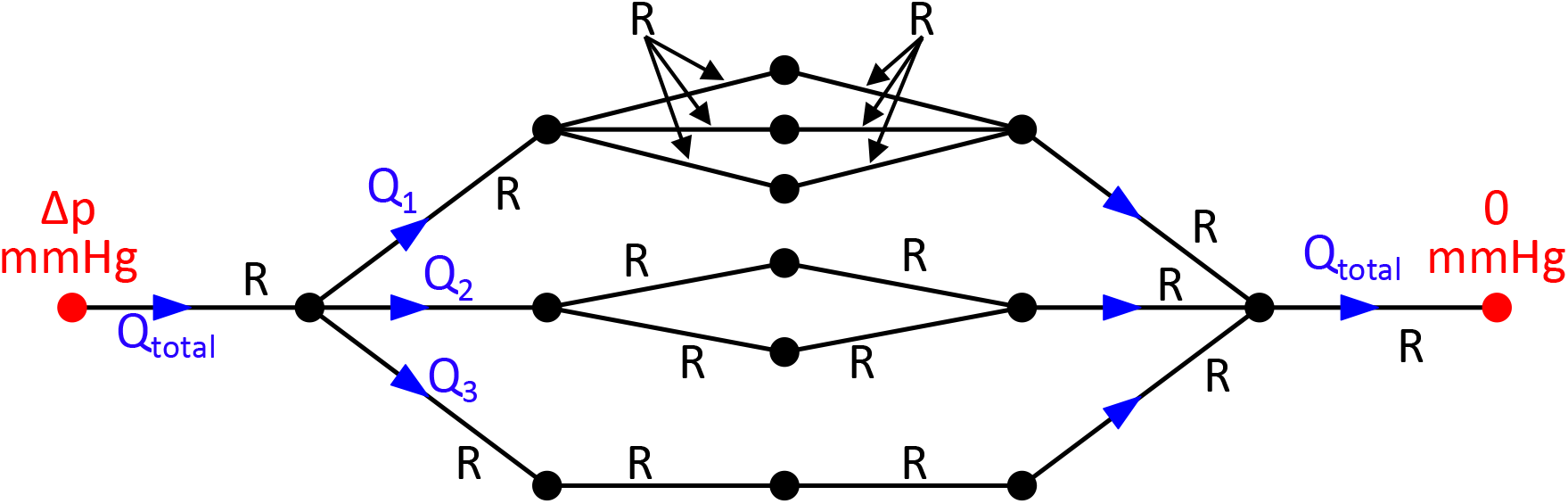
An analytical example illustrating how variable branching affects volume flow rate. This idealized example, in which flow is driven by an overall Δp pressure drop and every vessel segment has resistance R, can be solved analytically to find that 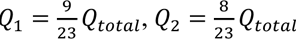, and 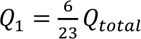 (i.e., branches with more numerous parallel channels draw larger amounts of flow due to decreased resistance). This effect explains the increased variability in volume flow rate, flow speed, and wall shear stress observed in the simulations with variable branching, compared to the cases with ideal trifurcations (Fig. 2).

**Fig. S3:**
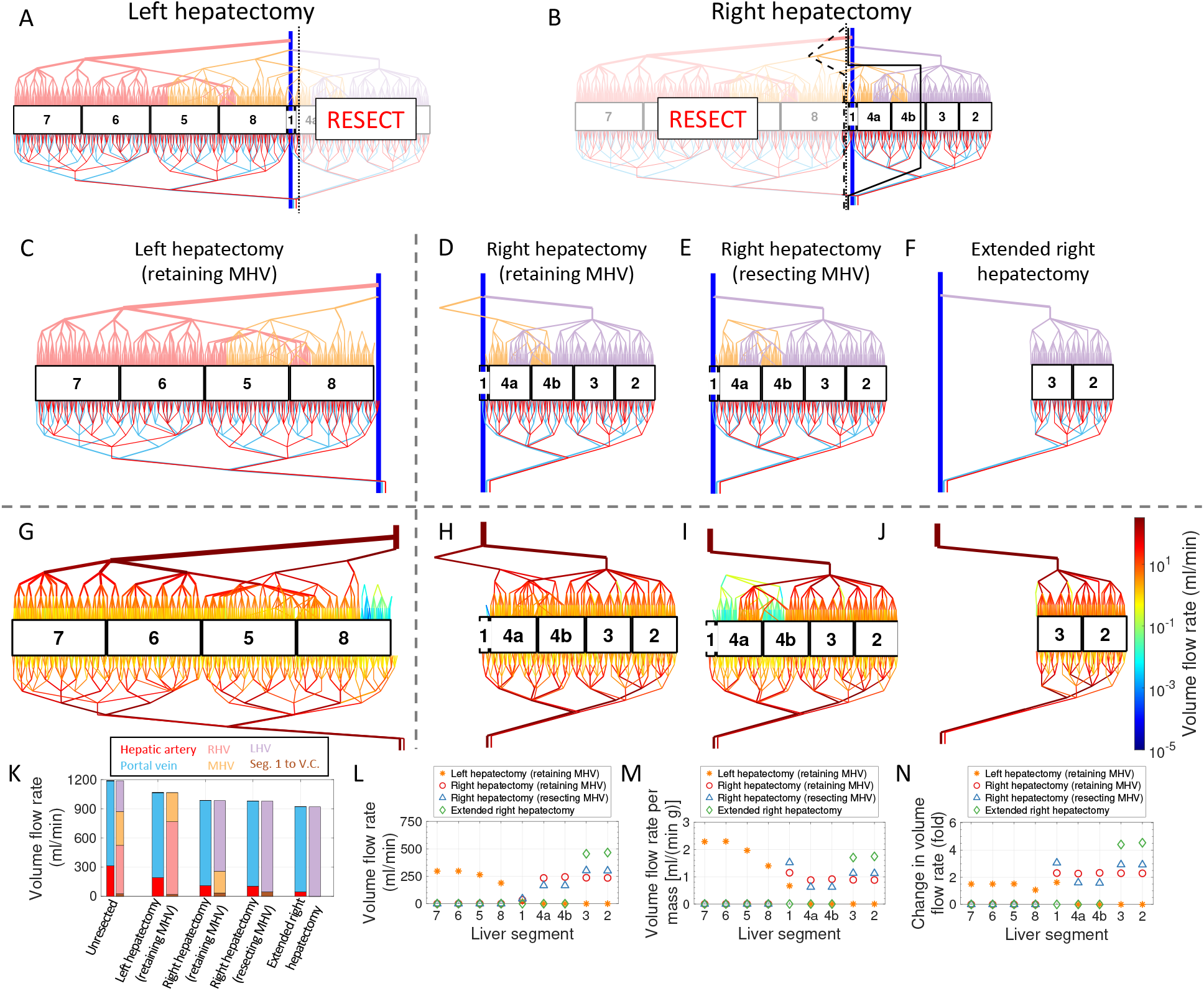
Simulations of four resections for an additional randomly generated network with variable branching. (A-B) Schematics of a left or right hepatectomy applied to a realistically sized network (containing 5 million lobules); dashed/dotted/solid black lines indicate potential surgical decisions. (C-F) Schematic illustration of networks following (C) left hepatectomy (33% resection) or (D-F) three variations of a right hepatectomy (67%, 67%, and 84% resection, respectively), as indicated. (G-J) Schematic plots with color encoding the volume flow rate. The color bar at the far right corresponds to all four plots. (K-N) Plots of the (K-L) volume flow rate, (M) volume flow rate per unit mass, and (N) fractional change in volume flow rate for different (K) vessels and (L-N) liver segments under all four resection scenarios. This figure is analogous to Fig. 3, but for a different network.

**Fig. S4:**
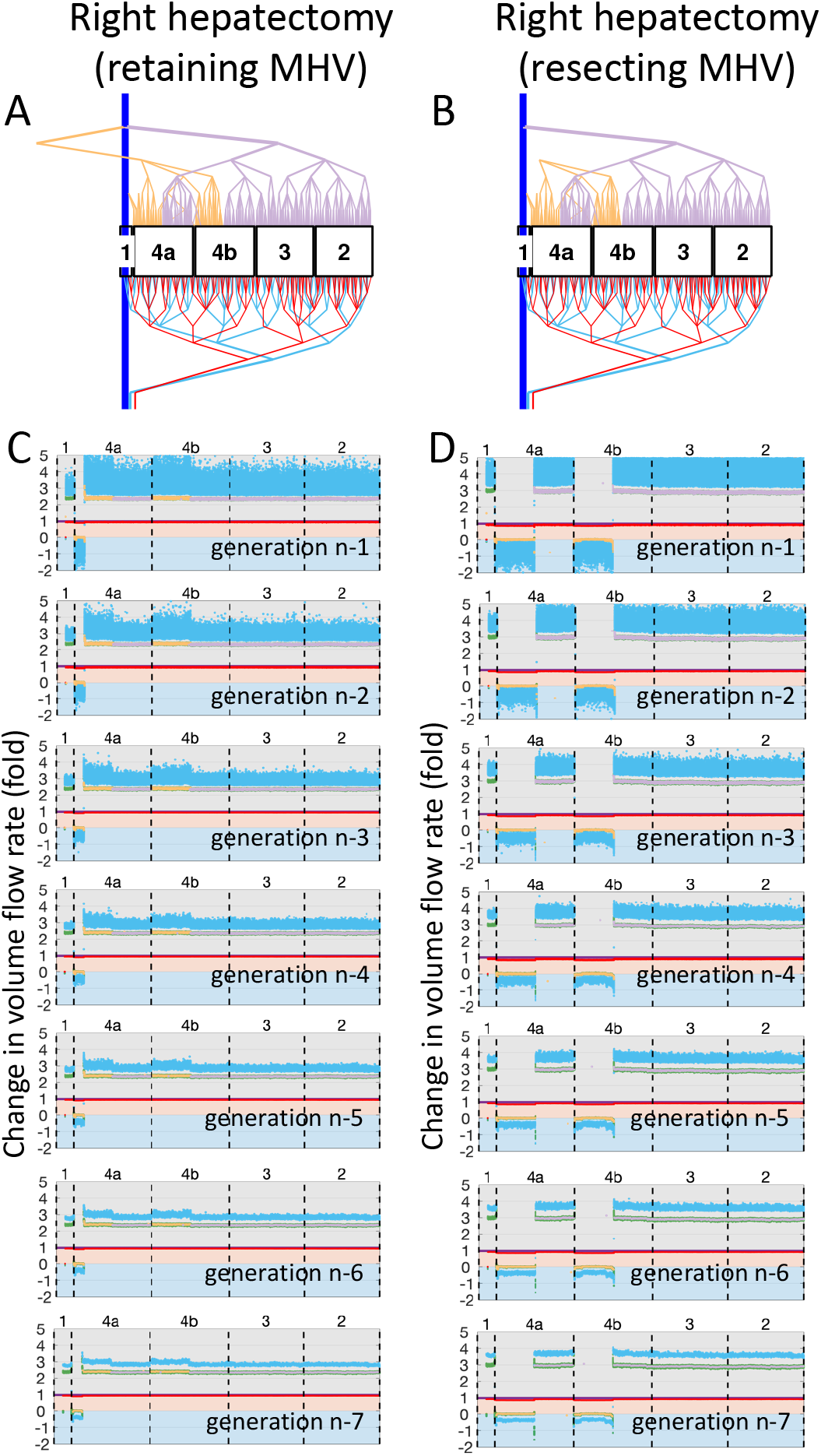
Increased volume flow rate and retrograde portal flow are persistent across several vascular generations. (A-B) Schematics of right hepatectomy with MHV (A) retention and (B) resection. Each figure indicates the applicable scenario for the plots in that same column. (C-D) Plots of the factor by which volume flow rate changes in the resected liver, compared to the unresected liver, for each color-coded vessel in each labeled liver segment following right hepatectomy with MHV (C) retention or (D) resection. Black labels indicate the generation number, where n corresponds to the sinusoids. Note that the corresponding unresected liver has 5 million lobules.

**Table S1:**
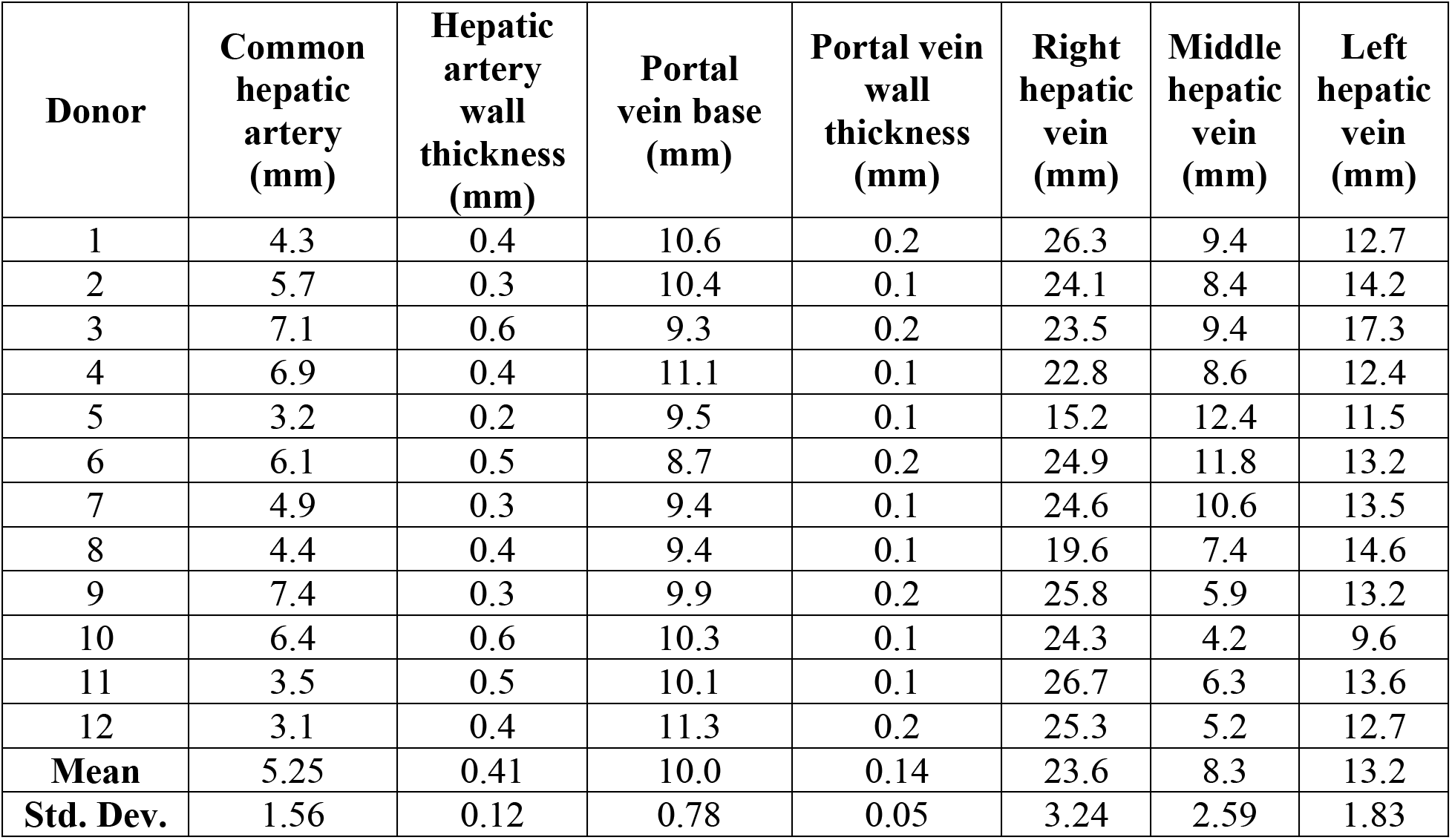
Vessel base diameter measurements from deceased liver donors. See Methods for details of how the lumen radii were calculated.

| **Donor** | **Common hepatic artery (mm)** | **Hepatic artery wall thickness (mm)** | **Portal vein base (mm)** | **Portal vein wall thickness (mm)** | **Right hepatic vein (mm)** | **Middle hepatic vein (mm)** | **Left hepatic vein (mm)** |
| --- | --- | --- | --- | --- | --- | --- | --- |
| 1 | 4.3 | 0.4 | 10.6 | 0.2 | 26.3 | 9.4 | 12.7 |
| 2 | 5.7 | 0.3 | 10.4 | 0.1 | 24.1 | 8.4 | 14.2 |
| 3 | 7.1 | 0.6 | 9.3 | 0.2 | 23.5 | 9.4 | 17.3 |
| 4 | 6.9 | 0.4 | 11.1 | 0.1 | 22.8 | 8.6 | 12.4 |
| 5 | 3.2 | 0.2 | 9.5 | 0.1 | 15.2 | 12.4 | 11.5 |
| 6 | 6.1 | 0.5 | 8.7 | 0.2 | 24.9 | 11.8 | 13.2 |
| 7 | 4.9 | 0.3 | 9.4 | 0.1 | 24.6 | 10.6 | 13.5 |
| 8 | 4.4 | 0.4 | 9.4 | 0.1 | 19.6 | 7.4 | 14.6 |
| 9 | 7.4 | 0.3 | 9.9 | 0.2 | 25.8 | 5.9 | 13.2 |
| 10 | 6.4 | 0.6 | 10.3 | 0.1 | 24.3 | 4.2 | 9.6 |
| 11 | 3.5 | 0.5 | 10.1 | 0.1 | 26.7 | 6.3 | 13.6 |
| 12 | 3.1 | 0.4 | 11.3 | 0.2 | 25.3 | 5.2 | 12.7 |
| **Mean** | 5.25 | 0.41 | 10.0 | 0.14 | 23.6 | 8.3 | 13.2 |
| **Std. Dev.** | 1.56 | 0.12 | 0.78 | 0.05 | 3.24 | 2.59 | 1.83 |

**Table S2:**
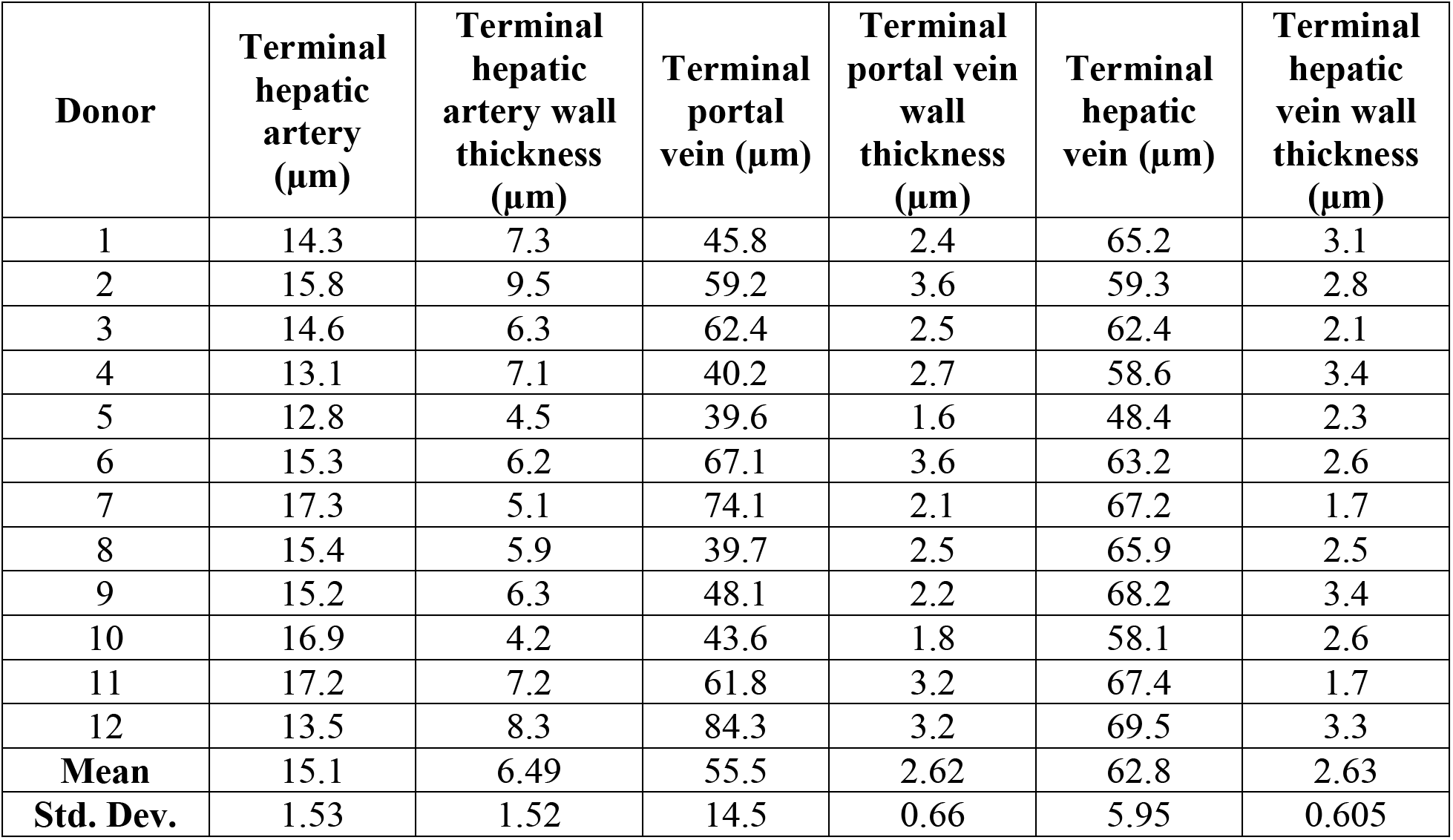
Terminal vessel diameter measurements (i.e., diameter measurements at the sinusoids) from deceased liver donors. See Methods for details of how the lumen radii were calculated.

## References

1. Lautt WW. Colloquium Series on Integrated Systems Physiology: From Molecule to Function to Disease. Hepatic Circulation: Physiology and Pathophysiology. San Rafael (CA): Morgan & Claypool Life Sciences Copyright © 2010 by Morgan & Claypool Life Sciences.; 2009.

2. Couinaud C. [The anatomy of the liver]. Ann Ital Chir. 1992;63(6):693–7.

3. Mr. Kiernan on the Anatomy and Physiology of the Liver. Med Chir Rev. 1834;20(40):305–15.

4. Teutsch HF, Schuerfeld D, Groezinger E. Three-dimensional reconstruction of parenchymal units in the liver of the rat. Hepatology. 1999;29(2):494–505.

5. Dahmen U, Hall CA, Madrahimov N, Milekhin V, Dirsch O. Regulation of hepatic microcirculation in stepwise liver resection. Acta Gastroenterol Belg. 2007;70(4):345–51.

6. van Mierlo KM, Schaap FG, Dejong CH, Olde Damink SW. Liver resection for cancer: New developments in prediction, prevention and management of postresectional liver failure. J Hepatol. 2016;65(6):1217–31.

7. Mise Y, Sakamoto Y, Ishizawa T, Kaneko J, Aoki T, Hasegawa K, et al. A worldwide survey of the current daily practice in liver surgery. Liver Cancer. 2013;2(1):55–66.

8. Makuuchi M, Thai BL, Takayasu K, Takayama T, Kosuge T, Gunvén P, et al. Preoperative portal embolization to increase safety of major hepatectomy for hilar bile duct carcinoma: a preliminary report. Surgery. 1990;107(5):521–7.

9. Deal R, Frederiks C, Williams L, Olthof PB, Dirscherl K, Keutgen X, et al. Rapid Liver Hypertrophy After Portal Vein Occlusion Correlates with the Degree of Collateralization Between Lobes-a Study in Pigs. J Gastrointest Surg. 2018;22(2):203–13.

10. Schnitzbauer AA, Lang SA, Goessmann H, Nadalin S, Baumgart J, Farkas SA, et al. Right portal vein ligation combined with in situ splitting induces rapid left lateral liver lobe hypertrophy enabling 2-staged extended right hepatic resection in small-for-size settings. Ann Surg. 2012;255(3):405–14.

11. Aragon RJ, Solomon NL. Techniques of hepatic resection. J Gastrointest Oncol. 2012;3(1):28–40.

12. Goja S, Kumar Yadav S, Singh Soin A. Readdressing the Middle Hepatic Vein in Right Lobe Liver Donation: Triangle of Safety. Liver Transpl. 2018;24(10):1363–76.

13. Guo HJ, Wang K, Chen KC, Liu ZK, Al-Ameri A, Shen Y, et al. Middle hepatic vein reconstruction in adult right lobe living donor liver transplantation improves recipient survival. Hepatobiliary Pancreat Dis Int. 2019;18(2):125–31.

14. Lu H, Wu L, Yuan R, Liao W, Lei J, Shao J. Modified median hepatic fissure approach for resection of liver tumours located in the angle between the root of the middle and right hepatic veins. BMC Surg. 2021;21(1):410.

15. Abdalla EK, Denys A, Chevalier P, Nemr RA, Vauthey JN. Total and segmental liver volume variations: implications for liver surgery. Surgery. 2004;135(4):404–10.

16. Araki K, Harimoto N, Kubo N, Watanabe A, Igarashi T, Tsukagoshi M, et al. Functional remnant liver volumetry using Gd-EOB-DTPA-enhanced magnetic resonance imaging (MRI) predicts post-hepatectomy liver failure in resection of more than one segment. HPB (Oxford). 2020;22(2):318–27.

17. Christ B, Collatz M, Dahmen U, Herrmann KH, Höpfl S, König M, et al. Hepatectomy-Induced Alterations in Hepatic Perfusion and Function - Toward Multi-Scale Computational Modeling for a Better Prediction of Post-hepatectomy Liver Function. Front Physiol. 2021;12:733868.

18. Emond JC, Goodrich NP, Pomposelli JJ, Baker TB, Humar A, Grant DR, et al. Hepatic Hemodynamics and Portal Flow Modulation: The A2ALL Experience. Transplantation. 2017;101(10):2375–84.

19. Paulsen AW, Klintmalm GB. Direct measurement of hepatic blood flow in native and transplanted organs, with accompanying systemic hemodynamics. Hepatology. 1992;16(1):100–11.

20. Pravisani R, Soyama A, Takatsuki M, Hidaka M, Adachi T, Ono S, et al. Relationship Between Venous Drainage Patterns and Regeneration of Segments 5 and 8 in Right Lobe Grafts in Adult Living-Donor Liver Transplant Recipients. Exp Clin Transplant. 2019;17(4):529–35.

21. Sureka B, Sharma N, Khera PS, Garg PK, Yadav T. Hepatic vein variations in 500 patients: surgical and radiological significance. Br J Radiol. 2019;92(1102):20190487.

22. Boissier N, Drasdo D, Vignon-Clementel IE. Simulation of a detoxifying organ function: Focus on hemodynamics modeling and convection-reaction numerical simulation in microcirculatory networks. Int J Numer Method Biomed Eng. 2021;37(2):e3422.

23. Debbaut C, Monbaliu D, Casteleyn C, Cornillie P, Van Loo D, Masschaele B, et al. From vascular corrosion cast to electrical analog model for the study of human liver hemodynamics and perfusion. IEEE Trans Biomed Eng. 2011;58(1):25–35.

24. van der Plaats A, Hart NA, Verkerke GJ, Leuvenink HG, Verdonck P, Ploeg RJ, et al. Numerical simulation of the hepatic circulation. Int J Artif Organs. 2004;27(3):222-30.

25. Audebert C, Bekheit M, Bucur P, Vibert E, Vignon-Clementel IE. Partial hepatectomy hemodynamics changes: Experimental data explained by closed-loop lumped modeling. J Biomech. 2017;50:202–8.

26. Golse N, Joly F, Combari P, Lewin M, Nicolas Q, Audebert C, et al. Predicting the risk of post-hepatectomy portal hypertension using a digital twin: A clinical proof of concept. J Hepatol. 2021;74(3):661–9.

27. Torres Rojas AM, Lorente S, Hautefeuille M, Sanchez-Cedillo A. Hierarchical Modeling of the Liver Vascular System. Front Physiol. 2021;12:733165.

28. Verma A MA, Melunis J, Hengstler JG, Vadigepalli R. From seeing to simulating: A survey of imaging techniques and spatially-resolved data for developing multiscale computational models of liver regeneration. Frontiers in Systems Biology. 2022;2.

29. Teutsch HF. The modular microarchitecture of human liver. Hepatology. 2005;42(2):317–25.

30. Ishikawa J, Takeo M, Iwadate A, Koya J, Kihira M, Oshima M, et al. Mechanical homeostasis of liver sinusoid is involved in the initiation and termination of liver regeneration. Commun Biol. 2021;4(1):409.

31. Cabrera M FU. Novel in vivo imaging techniques for the liver microvasculature. IntraVital. 2012;1(2):107–14.

32. Ghallab A HJ. Liver regeneration and new technical possibilities by two-photon based intravital imaging. SVU-International Journal of Veterinary Sciences. 2018;1(1):4–15.

33. Reif R, Ghallab A, Beattie L, Günther G, Kuepfer L, Kaye PM, et al. In vivo imaging of systemic transport and elimination of xenobiotics and endogenous molecules in mice. Arch Toxicol. 2017;91(3):1335–52.

34. Bonfiglio A, Leungchavaphongse K, Repetto R, Siggers JH. Mathematical modeling of the circulation in the liver lobule. J Biomech Eng. 2010;132(11):111011.

35. J Hu SLu, S Feng, M Long. Flow dynamics analyses of pathophysiological liver lobules using porous media theory. Acta Mechanica Sinica. 2017;33(4):823–32.

36. Lorente S, Hautefeuille M, Sanchez-Cedillo A. The liver, a functionalized vascular structure. Sci Rep. 2020;10(1):16194.

37. Ricken T, Dahmen U, Dirsch O. A biphasic model for sinusoidal liver perfusion remodeling after outflow obstruction. Biomech Model Mechanobiol. 2010;9(4):435–50.

38. Blumgart’s Surgery of the Liver, Biliary Tree and Pancreas, 2-volume set, Elsevier, 2017.

39. Hashimoto K, Murakami T, Dono K, Hori M, Kim T, Kudo M, et al. Assessment of the severity of liver disease and fibrotic change: the usefulness of hepatic CT perfusion imaging. Oncol Rep. 2006;16(4):677–83.

40. Radtke A, Nadalin S, Sotiropoulos GC, Molmenti EP, Schroeder T, Valentin-Gamazo C, et al. Computer-assisted operative planning in adult living donor liver transplantation: a new way to resolve the dilemma of the middle hepatic vein. World J Surg. 2007;31(1):175–85.

41. Sato Y, Koyama S, Tsukada K, Hatakeyama K. Acute portal hypertension reflecting shear stress as a trigger of liver regeneration following partial hepatectomy. Surg Today. 1997;27(6):518–26.

42. Chouillard EK, Gumbs AA, Cherqui D. Vascular clamping in liver surgery: physiology, indications and techniques. Ann Surg Innov Res. 2010;4:2.

43. Guglielmi A, Ruzzenente A, Conci S, Valdegamberi A, Iacono C. How much remnant is enough in liver resection? Dig Surg. 2012;29(1):6–17.

44. Khan AS, Garcia-Aroz S, Ansari MA, Atiq SM, Senter-Zapata M, Fowler K, et al. Assessment and optimization of liver volume before major hepatic resection: Current guidelines and a narrative review. Int J Surg. 2018;52:74–81.

45. Lautt WW. Mechanism and role of intrinsic regulation of hepatic arterial blood flow: hepatic arterial buffer response. Am J Physiol. 1985;249(5 Pt 1):G549-56.

46. Benoit JN, Womack WA, Hernandez L, Granger DN. “Forward” and “backward” flow mechanisms of portal hypertension. Relative contributions in the rat model of portal vein stenosis. Gastroenterology. 1985;89(5):1092–6.

47. Denys AL, Abehsera M, Leloutre B, Sauvanet A, Vilgrain V, O’Toole D, et al. Intrahepatic hemodynamic changes following portal vein embolization: a prospective Doppler study. Eur Radiol. 2000;10(11):1703–7.

48. Karanjia ND, Lordan JT, Quiney N, Fawcett WJ, Worthington TR, Remington J. A comparison of right and extended right hepatectomy with all other hepatic resections for colorectal liver metastases: a ten-year study. Eur J Surg Oncol. 2009;35(1):65–70.

49. Emond JC, Renz JF, Ferrell LD, Rosenthal P, Lim RC, Roberts JP, et al. Functional analysis of grafts from living donors. Implications for the treatment of older recipients. Ann Surg. 1996;224(4):544–52; discussion 52-4.

50. Cattral MS, Molinari M, Vollmer CM, Jr., McGilvray I, Wei A, Walsh M, et al. Living-donor right hepatectomy with or without inclusion of middle hepatic vein: comparison of morbidity and outcome in 56 patients. Am J Transplant. 2004;4(5):751–7.

51. Nobuoka T, Mizuguchi T, Oshima H, Shibata T, Kimura Y, Mitaka T, et al. Portal blood flow regulates volume recovery of the rat liver after partial hepatectomy: molecular evaluation. Eur Surg Res. 2006;38(6):522–32.

52. Schoen JM, Wang HH, Minuk GY, Lautt WW. Shear stress-induced nitric oxide release triggers the liver regeneration cascade. Nitric Oxide. 2001;5(5):453–64.

53. Francavilla A, Zeng Q, Polimeno L, Carr BI, Sun D, Porter KA, et al. Small-for-size liver transplanted into larger recipient: a model of hepatic regeneration. Hepatology. 1994;19(1):210–6.

54. Mars WM, Liu ML, Kitson RP, Goldfarb RH, Gabauer MK, Michalopoulos GK. Immediate early detection of urokinase receptor after partial hepatectomy and its implications for initiation of liver regeneration. Hepatology. 1995;21(6):1695–701.

55. Taub R. Liver regeneration 4: transcriptional control of liver regeneration. Faseb j. 1996;10(4):413–27.

56. Fausto N, Laird AD, Webber EM. Liver regeneration. 2. Role of growth factors and cytokines in hepatic regeneration. Faseb j. 1995;9(15):1527–36.

57. Chen F, Jimenez RJ, Sharma K, Luu HY, Hsu BY, Ravindranathan A, et al. Broad Distribution of Hepatocyte Proliferation in Liver Homeostasis and Regeneration. Cell Stem Cell. 2020;26(1):27–33.e4.

58. Dirsch O, Madrahimov N, Chaudri N, Deng M, Madrahimova F, Schenk A, et al. Recovery of liver perfusion after focal outflow obstruction and liver resection. Transplantation. 2008;85(5):748–56.

59. Panis Y, McMullan DM, Emond JC. Progressive necrosis after hepatectomy and the pathophysiology of liver failure after massive resection. Surgery. 1997;121(2):142–9.

60. Allard MA, Adam R, Bucur PO, Termos S, Cunha AS, Bismuth H, et al. Posthepatectomy portal vein pressure predicts liver failure and mortality after major liver resection on noncirrhotic liver. Ann Surg. 2013;258(5):822-9; discussion 9-30.

61. Kawai M, Naruse K, Komatsu S, Kobayashi S, Nagino M, Nimura Y, et al. Mechanical stress-dependent secretion of interleukin 6 by endothelial cells after portal vein embolization: clinical and experimental studies. J Hepatol. 2002;37(2):240–6.

62. Ikegami T, Yoshizumi T, Soejima Y, Ikeda T, Kawanaka H, Uchiyama H, et al. Application of splenectomy to decompress portal pressure in left lobe living donor liver transplantation. Fukuoka Igaku Zasshi. 2013;104(9):282–9.

63. Winslow ER, Brunt LM. Perioperative outcomes of laparoscopic versus open splenectomy: a meta-analysis with an emphasis on complications. Surgery. 2003;134(4):647–53; discussion 54-5.

64. Watanabe Y, Horiuchi A, Yoshida M, Yamamoto Y, Sugishita H, Kumagi T, et al. Significance of laparoscopic splenectomy in patients with hypersplenism. World J Surg. 2007;31(3):549–55.

65. Ikegami T, Shirabe K, Soejima Y, Yoshizumi T, Uchiyama H, Yamashita Y, et al. Strategies for successful left-lobe living donor liver transplantation in 250 consecutive adult cases in a single center. J Am Coll Surg. 2013;216(3):353–62.

66. Condon RE, Chapman ND, Nyhus LM, Harkins HN. Hepatic arterial and portal venous pressure-flow relationships in isolated, perfused liver. Am J Physiol. 1962;202:1090–4.

67. Hanson KM. Dilator responses of the canine hepatic vasculature. Angiologica. 1973;10(1):15–23.

68. Hanson KM, Johnson PC. Local control of hepatic arterial and portal venous flow in the dog. Am J Physiol. 1966;211(3):712–20.

69. Messmer K, Brendel W, Devens K, Reulen HJ. [The pressure relationship of hepatic circulation]. Pflugers Arch Gesamte Physiol Menschen Tiere. 1966;289(1):75–90.

70. Torrance HB. The control of the hepatic arterial circulation. J Physiol. 1961;158(1):39–49.

71. Jakab F, Ráth Z, Schmal F, Nagy P, Faller J. The interaction between hepatic arterial and portal venous blood flows; simultaneous measurement by transit time ultrasonic volume flowmetry. Hepatogastroenterology. 1995;42(1):18–21.

72. Lautt WW, Legare DJ, Ezzat WR. Quantitation of the hepatic arterial buffer response to graded changes in portal blood flow. Gastroenterology. 1990;98(4):1024–8.

73. Sato Y, Tsukada K, Hatakeyama K. Role of shear stress and immune responses in liver regeneration after a partial hepatectomy. Surg Today. 1999;29(1):1–9.

74. Marcos A, Olzinski AT, Ham JM, Fisher RA, Posner MP. The interrelationship between portal and arterial blood flow after adult to adult living donor liver transplantation. Transplantation. 2000;70(12):1697–703.

75. Jessen E, Steinbach MC, Debbaut C, Schillinger D. Rigorous mathematical optimization of synthetic hepatic vascular trees. J R Soc Interface. 2022;19(191):20220087.

76. Pries AR, Secomb TW. Microvascular blood viscosity in vivo and the endothelial surface layer. Am J Physiol Heart Circ Physiol. 2005;289(6):H2657–64.

77. Oechtering TH, Roberts GS, Panagiotopoulos N, Wieben O, Reeder SB, Roldán-Alzate A. Clinical Applications of 4D Flow MRI in the Portal Venous System. Magn Reson Med Sci. 2022;21(2):340–53.

78. Rutkowski DR, Reeder SB, Fernandez LA, Roldán-Alzate A. Surgical planning for living donor liver transplant using 4D flow MRI, computational fluid dynamics and in vitro experiments. Comput Methods Biomech Biomed Eng Imaging Vis. 2018;6(5):545–55.

79. Hyodo R, Takehara Y, Naganawa S. 4D Flow MRI in the portal venous system: imaging and analysis methods, and clinical applications. Radiol Med. 2022;127(11):1181–98.

80. Takahashi H, Shigefuku R, Yoshida Y, Ikeda H, Matsunaga K, Matsumoto N, et al. Correlation between hepatic blood flow and liver function in alcoholic liver cirrhosis. World J Gastroenterol. 2014;20(45):17065–74.

81. Thng CH, Koh TS, Collins DJ, Koh DM. Perfusion magnetic resonance imaging of the liver. World J Gastroenterol. 2010;16(13):1598–609.

82. Byk K, Jasinski K, Bartel Z, Jasztal A, Sitek B, Tomanek B, et al. MRI-based assessment of liver perfusion and hepatocyte injury in the murine model of acute hepatitis. Magma. 2016;29(6):789–98.

83. Balfour DC, Jr., Reynolds TB, Levinson DC, Mikkelsen WP, Pattison AC. Hepatic vein pressure studies for evaluation of intrahepatic portal hypertension. AMA Arch Surg. 1954;68(4):442-7.

84. Debbaut C, Segers P, Cornillie P, Casteleyn C, Dierick M, Laleman W, et al. Analyzing the human liver vascular architecture by combining vascular corrosion casting and micro-CT scanning: a feasibility study. J Anat. 2014;224(4):509–17.

85. Kline TL, Zamir M, Ritman EL. Relating function to branching geometry: a micro-CT study of the hepatic artery, portal vein, and biliary tree. Cells Tissues Organs. 2011;194(5):431–42.

